# Sex-specific deubiquitylation drives immune-related neurodegeneration in *Drosophila*

**DOI:** 10.1101/2022.12.09.519782

**Authors:** Jingnu Xia, Adán Pinto-Fernández, Andreas Damianou, Jeffery Y Lee, Benedikt M Kessler, Ilan Davis, Paul Elliott, Petros Ligoxygakis

**Author notes:** Joint corresponding authors and.

## Abstract

Risk of neurodegenerative disease such as late onset Alzheimer’s is linked to aberrant ubiquitinylation and accumulation of non-degraded proteins in brain cells. A glial network of innate immune genes modulates inflammatory responses to such protein deposition. However, vulnerability differs between the sexes. Here, we show that the *Drosophila* homologue of the deubiquitylase Trabid can align the sex-specific aspects of neurodegenerative phenotypes with changes in ubiquitylation and inflammatory activity. An enzymatically null Trabid in flies, caused sex-specific changes in locomotion, sleep patterns, brain histology and ultimately, lifespan. These changes were underscored by altered ubiquitin and proteome enrichment profiles and the same enzymatic activity as its human counterpart. When the sex-determination gene *transformer* was silenced in astrocytes or immunocompetent tissues, sex differences were significantly reduced. Our results indicate that Trabid underscores sex-specificity in disease neurology, by controlling the balance between ubiquitylation and inflammation.

## Introduction

Neurodegenerative diseases present different risks depending on sex assigned at birth. Alzheimer’s Disease (AD) is approx. 1.7 times more frequent in females than males (Rajan et al., 2021) whereas prevalence of Parkinson’s Disease (PD) is 1.5 times greater in males (Reekes et al., 2020) with a steeper increase in the 60–69 and 70–79 decades of life (Hirsch et al., 2016). Common symptoms include gait and locomotion impairment, sleep dysregulation and cognition decline (Bombois *et al,* 2010; Pieruccini-Faria *et al,* 2021). Studies have shown that the different risk ratio may be due to sex-specific lifespan, hormonal, genetic effects and reproductive factors (Erkkinen et al., 2018). However, identification of molecular drivers that underscore sex-specificity in neurodegenerative disease remains rare.

Ubiquitylation and deubiquitylation are dynamic posttranslational modifications that coordinate to modify a range of cellular processes including inflammation (Kulathu and Komander, 2012). The ubiquitylation process includes ubiquitin-activating enzyme (E1), ubiquitin conjugating enzyme (E2) and ubiquitin ligase (E3), which are the “writers”. E3s provide specificity to the polyubiquitin chain elongation and substrate recognition. After the signal is decoded and passed on, ubiquitin signals need to be “erased” by ubiquitin hydrolysis enzymes or deubiquitylases (DUBs). DUBs usually consist of a catalytic domain and a domain for substrate recognition and binding (Schmidt et al., 2021). Polymeric ubiquitin chains and DUBs control signalling downstream of innate immune receptors such as nuclear oligomerization domain (NOD)-like receptors (NLRs) and tumour necrosis factor (TNF) receptor 1 (TNFR1) to activate nuclear factor κB (NF-κB)-mediated gene activation (Damgaard et al., 2016; Emmerich et al., 2013; Gerlach et al., 2011; Kensche et al., 2016).

Aggregation of misfolded proteins, including amyloid β (Aβ), tau and α-synuclein, can progressively damage the nervous system and are considered hallmarks of AD and PD. It has been proposed that modulating the ubiquitylation level-via ligase or DUB activity-of these proteins could regulate accumulation (Ciechanover and Kwon, 2015). Reducing the E3 ligase HRD1-mediated protein degradation can cause the accumulation of Aβ and amyloid precursor protein (APP) (Kaneko et al., 2010, p. 1). The carboxyl terminus of heat-shock cognate HSP70-interacting protein (CHIP) is a E3 ligase that can regulate tau ubiquitylation and degradation in cell culture (Shin et al., 2005). Overexpressing CHIP could ameliorate the early development of tau in animal model (Sahara et al., 2005). Knocking out DUB Usp8 promotes the lysosomal degradation of α-synuclein in human cells, while knocking out Usp8 in *Drosophila*, protected the animal from locomotor defects and cell loss (Alexopoulou et al., 2016). Moreover, knocking out DUB USP9X caused the monoubiquitylation of α synuclein and reduced the efficiency of α-synuclein autophagy (Rott et al., 2011).

Dysregulation of degrading misfolded proteins is not the sole source of pathogenesis in neurodegenerative disease. The ubiquitin-like protein — Ubiquilin 1 affects the trafficking of Amyloid Precursor Protein and the generation of Aβ-amyloid (Hiltunen et al., 2006). Members of the NEDD4 family of E3 ligases are involved in PD not only in regulating the clearance of oligomeric α-synuclein but interacting with additional substrates and interacting proteins in affected tissues (reviewed in Conway et al., 2022). NEDD4 promotes α-synuclein degradation through the endosomal-lysosomal pathway (Tofaris et al., 2011) while inactive or reduced levels of NEDD4 can protect neurons from zinc-induced cell death and apoptosis (Kwak et al., 2012). In addition to aggregated α-synuclein, PD patients have elevated pro-inflammatory cytokine levels in their serums (Dzamko et al., 2015). Mutations in the E3 ligase Parkin and the ubiquitin kinase PINK1 indicate their involvement in familial forms of PD by regulating mitochondria homeostasis and inflammation (reviewed in Pickrell and Youle, 2015; see also Henn et al., 2007; Sliter et al., 2018). Loss of Parkin caused aged mice to develop neurodegenerative phenotypes through overactive immunity and STING/NF-κB-induced inflammation (Sliter et al., 2018). The link between constitutive inflammatory activity and neurodegeneration seems to be evolutionary conserved (reviewed in Arora and Ligoxygakis, 2020; Kaltschmidt et al., 2022). In *Drosophila,* flies genetically predisposed to high immune responses show age-dependent neurodegeneration (Kounatidis et al., 2017; Shukla et al., 2019). Silencing NF-κB in glia or neurons, blocked neurodegeneration and associated neurological phenotypes (Kounatidis et al., 2017; Shukla et al., 2019).

One of the regulators of NF-κB-mediated inflammation is the DUB Trabid (Fernando et al., 2014; Hua et al., 2022; Kounatidis et al., 2017). In *Drosophila,* Trabid acts as a negative regulator of the IMD/NF-κB pathway, homologous to TNFR1/NF-κB pathway (Fernando et al., 2014; Hua et al., 2022; Kounatidis et al., 2017). Deletion of *trabid* (*trbd*), de-represses the pathway and results in increased neuroinflammation, locomotion defects and ultimately brain lesions associated with age-dependent neurodegeneration (Kounatidis et al., 2017). Trabid is an A20-like family DUB, evolutionary conserved from flies to humans (Fernando et al., 2014; Tran et al., 2008; see also below). Human Trabid has been shown to preferentially hydrolyse Lys29, 33 and 63 diubiquitin chains *in vitro* (Licchesi et al., 2012). Trabid has also been reported as a positive regulator of the Wnt pathway in both mammalian cells and *Drosophila* flies (Tran et al., 2008). Moreover, Trabid knockdown promoted hepatocellular carcinoma growth through Twist1 ubiquitination (Zhu et al., 2019). Finally, hydrolysis of ubiquitin on histones by Trabid activated TLR-dependent cytokine release (Jin et al., 2016). Knocking out Trabid in mouse dendritic cells epigenetically suppressed Toll-like receptor (TLR)-mediated NF-κB signalling and associated inflammation in the central nervous system (CNS) (Jin et al., 2016). Taken together the results above show a role for Trabid in brain and CNS immunity in both flies and mice. However, how Trabid affects brain ubiquitin homeostasis, brain inflammation, development and behaviour in a sex specific manner remains unclear.

Here we show that *Drosophila* Trabid has the same enzymatic activity as its human counterpart. Introducing, through CRISPR-Cas9, an enzymatically *null* point mutation (*C518A*) in flies caused sex-specific changes inbehaviour and life expectancy. In assays of lifespan, locomotion, and sleep, the *C518A* mutation had more detrimental effects in the males than in the females. These sexually dimorphic changes were underscored by brain histology, inflammation marker expression, ubiquitin/SUMO enrichment and total proteome and ubiquitome analysis. Some of these sex-biased differences were reduced when the sex determination gene *transformer (tra)* was silenced in astrocytes or in immunocompetent tissues. Our results highlight the sex-specific impact of the DUB Trabid in age-dependent changes of neurological behaviours and brain function.

## Results

### The Ank-OTU enzymatic region in *Drosophila* Trabid preferentially cleaves Lys-29 and Lys-33 ubiquitin chains *in vitro*

Aligning the human, mouse, and *Drosophila* Trabid protein sequences indicated significant evolutionary conservation (Fig. 1A). Trabid has three NZF ubiquitin binding domains at the N terminus and an Ank-OTU domain, which provides the enzymatic activity, at the C terminus (Licchesi et al., 2012) (Fig. 1A). Superimposition of the human Trabid^AnkOTU^ structure (PDB:3ZRH) (Licchesi et al., 2012) with the *Drosophila* AlphaFold2 predicted Trabid^AnkOTU^ structure indicated that the two were highly conserved, suggesting that the fly Trabid^AnkOTU^ had similar enzymatic activity as its human counterpart (Fig. 1B). Previous studies have indicated that human Trabid^AnkOTU^ showed a preferred enzymatic activity towards Lys29 and Lys33 diubiquitin chains while less towards Lys63 diubiquitin chains (Licchesi et al., 2012). We used 150 nM purified dTrabid^AnkOTU^ to individually hydrolyse all eight types of diubiquitin chains (3.5 µM) for 0, 5 and 30 minutes at room temperature. We observed that dTrabid^AnkOTU^ had a predominant cleaving activity toward Lys29 and 33 diubiquitin chains (Fig. 1C). We did not observe the hydrolysis of Lys63 chains at 150 nM DUB concentration (Fig. 1C), but increasing the DUB concentration to 300 nM, resulted in the hydrolysis of Lys63 diubiquitin (Fig. 1D, see below).

**Figure 1.**
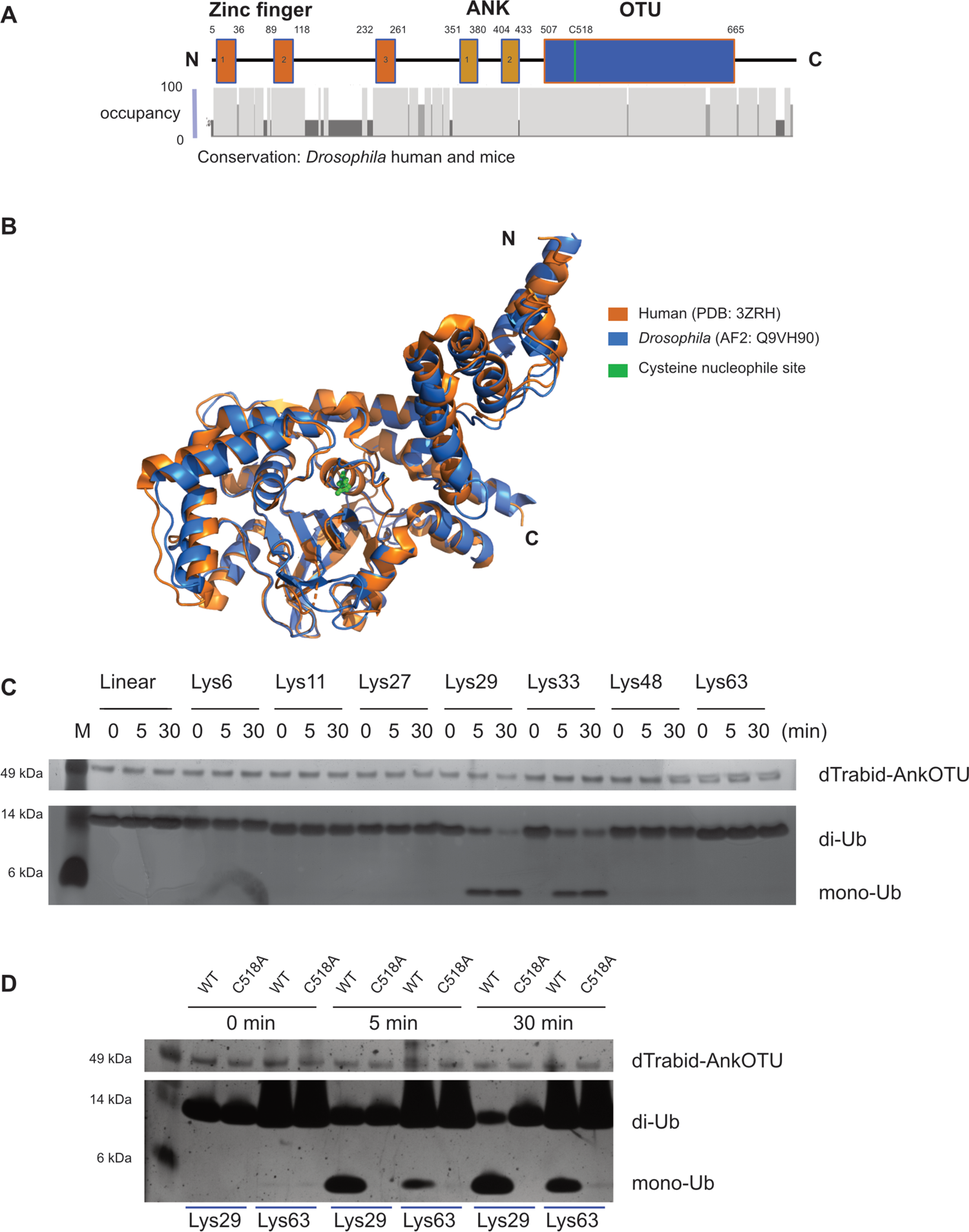
Comparison of *Drosophila* Trabid (dTrabid) with its human homologue (hTrabid) at the level of primary sequence, 3D structures and enzymatic activity. **(A)** Alignment of *Drosophila* (Q9VH90), human (Q9UGI0) and mice (Q7M760) Trabid amino acid sequence and annotated domain and nucleophile site of dTrabid. **(B)** Structure alignment of hTrabid-AnkOTU domain (3ZRH) and AlphaFold2 predicted dTrabid-AnkOTU (Q9VH90). **(C)** DUB assay of 150 nM dTrabid-AnkOTU on 3.5 μM eight different diubiquitin chain for 0, 5 and 30 minutes. dTrabid-AnkOTU preferentially hydrolyses Lys29 and Lys33 diubiquitin chain. **(D)** DUB assay of 300 nM dTrabid-AnkOTU (WT) and dTrabid-AnkOTUC518A (C518A) on 3.5 μM Lys29 and 63 diubiquitin chain for 0, 5 and 30 minutes. WT has a relative lower enzymatic activity towards Lys63 diubiquitin chain than Lys29 diubiquitin chain. The C518A mutation abrogates the enzymatic activity of Trabid.

Mutating the nucleophilic site (C443S) of human Trabid, resulted in loss of the enzyme’s hydrolytic activity (Licchesi et al., 2012). In *Drosophila* Trabid, the equivalent site was predicted to be Cysteine 518 (C518). We introduced a change to Alanine at that site (C518A) in the dTrabid^AnkOTU^ construct (dTrabid^AnkOTUC518A^) to assess its enzymatic activity towards Lys29 and Lys63 diubiquitin chains. 300 nM of dTrabid^AnkOTU^ or dTrabid^AnkOTUC518A^ were used to hydrolyse 3.5 µM Lys29 and Lys63 diubiquitin chains for 0, 5 and 30 minutes (Fig. 1D). As expected from experiments shown in Fig 1C also in this context, the wild-type enzyme significantly cleaved Lys29 and Lys63 diubiquitin (Fig. 1D). In contrast, limited activity towards both diubiquitin chains was observed for the mutant dTrabid^AnkOTUC518A^ (Fig. 1D). Hence, identical to its human counterpart (Licchesi et al., 2012), dTrabid^AnkOTU^ had a dual specificity against Lys29 and Lys33 ubiquitin chains and was less active towards Lys63 chains.

### Lifespan reduction in *trbd^C518A^* flies is sex-specific

To understand the effects of inactivating Trabid *in vivo,* we generated *trbd^C518A^* flies via CRISPR-Cas9 gene editing in the *w^1118^* background and homozygous lines were checked by Sanger sequencing. Introducing *trbd^C518A^,* resulted in significant reductions in lifespan in both males and females (p<0.0001 and p<0.01). Compared to the genetic background, the female mutant was affected less than the male in their survival probability (p<0.001) (Fig. 2A). Of note, that there was no difference in survival between the *w^1118^* females and males (Fig. 2A). This suggested that inactivated Trabid significantly affected the *Drosophila* lifespan in a sex-specific manner.

**Figure 2.**
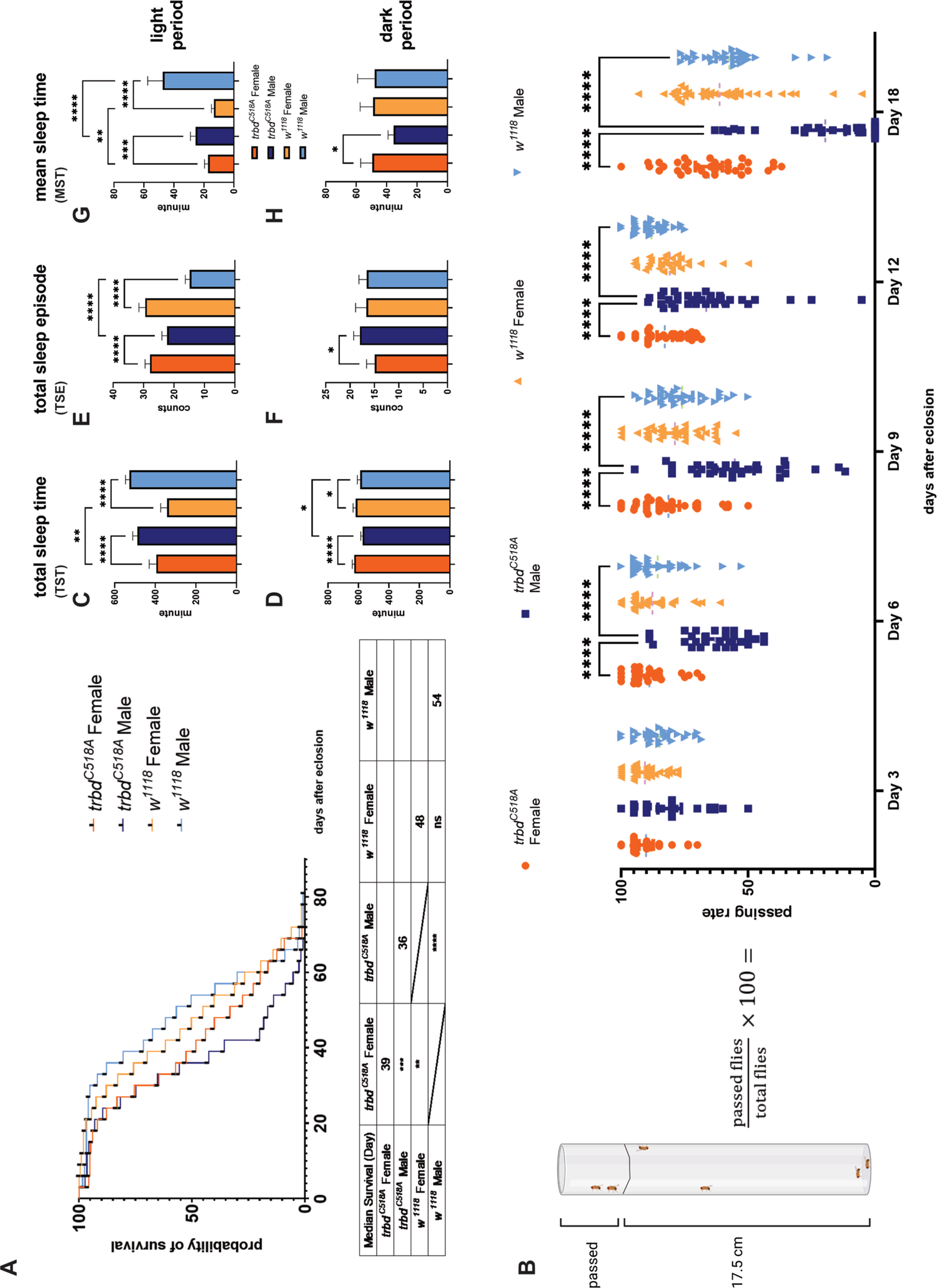
Lifespan, climbing, and sleep assays show that the *trbd^C518A^* mutation is more deleterious to male flies. **(A)** Survival plot of *trbd^C518A^* and *w^1118^* in females and males (log rank test, n=∼150). No significant difference between *w^1118^* female and male lifespan was observed. Male *trbd^C518A^* flies had a reduced lifespan compared to female *trbd^C518A^* (***p<0.001) and male *w^1118^* controls (****p<0.0001). Female *trbd^C518A^* had a reduced lifespan compared to *w^1118^* female controls (**p<0.01). **(B)** Climbing assay to test locomotor activity of *trbd^C518A^* and *w^1118^* at days 3, 6, 9, 12 and 18 post eclosion (two-way ANOVA with mixed model of interaction effect of time and genotype and Tukey HSD test, n=∼20 flies x 30 tests, 95% CI). There was no significant difference detected at day 3. Male *trbd^C518A^* flies had significantly reduced climbing passing rate compared to *trbd^C518A^* females and *w^1118^* control males at days 6, 9, 12 and 18 (all ****p<0.0001). (C-H) DAM result for total sleep time (TST), total sleep episode (TSE) and mean sleep time (MST) at light period (LP) and dark period (DP) (one-way ANOVA Kruskal-Wallis test with uncorrected Dunn’s test, n=∼66, 95% CI). **(C)** Males had more TST than females for both *trbd^C518A^* and *w^1118^* (both ****p<0.0001) and *trbd^C518A^* females had increased TST compared to *w^1118^* females in LP (**p<0.01). **(D)** Females had more TST than males for both *trbd^C518A^* (****p<0.0001) and *w^1118^* (*p<0.05) and *trbd^C518A^* males had reduced TST than *w^1118^* males (*p<0.05), in DP. **(E)** Females had significantly increased TSE compared to males in both *trbd^C518A^* and *w^1118^* (both ****p<0.0001) and *trbd^C518A^* males had significantly increased TSE compared to *w^1118^* male (****p<0.0001) in LP. **(F)** In DP, *trbd^C518A^* males had significant reduced TSE compared to *trbd^C518A^* females (*p<0.05). **(G)** In LP, females had significant less MST than males in both *trbd^C518A^* (***p<0.001) and *w^1118^* (****p<0.0001). *trbd^C518A^* female had increased MST compared to *w^1118^* female (**p<0.01) and *trbd^C518A^* males had decreased MST compared to *w^1118^* males (****p<0.0001). **(H)** In DP, *trbd^C518A^* females had increased MST compared to *w^1118^* females (*p<0.05).

### *trbd^C518A^* results in sex-specific locomotor defects in early adulthood

Locomotor defects are symptoms of and are used as assessment assays in neurodegenerative diseases (Pieruccini-Faria et al., 2021). In *Drosophila,* climbing assays have been generally used for evaluation of locomotion. Here we adapted an assay that can identify locomotion defects in early adulthood (Madabattula et al., 2015). We assessed our flies at day 3, 6, 9, 12 and 18 after eclosion. Flies passing the 17.5 cm mark on the graduate cylinder were considered as a pass (Fig. 2B). We found that *trbd^C518A^* males had a lower climbing passing rate than *w^1118^* males at day 6, 9, 12 and 18 (p<0.0001), while there was no significant difference between mutant and wild type females. We recorded that *trbd^C518A^* females had a higher pass rate throughout all timepoints compared to the *trbd^C518A^* males (p<0.0001), while no significant difference in climbing was recorded between *w^1118^* males and *w^1118^* females. This demonstrated that *trbd^C518A^* contributed to locomotion defects, but this effect was limited to male flies.

### Fragmented sleep in *trbd^C518A^* male flies

Sleep deficits have been closely associated with neurodegenerative diseases and AD patients have notable changes in their slow wave sleep before any cognitive decline diagnosis (Mander et al., 2015; reviewed in Wang and Holtzman, 2020). *Drosophila* and humans share critical sleep/wake circuits (reviewed in Cirelli and Bushey, 2008), and sleep disruption has been recorded for some *Drosophila* models of neurodegeneration (Buhl et al., 2019; Dissel, 2020; Luna et al., 2017, p.; Tabuchi et al., 2015). Unlike humans however, flies have a continuous sleep-wake cycle throughout the day, with a different pattern between daytime and night (Cirelli and Bushey, 2008). Here, we conducted sleep analysis in the light period (LP) and dark period (DP) of *trbd^C518A^* mutants and their *w^1118^* genetic background. The sleep pattern characteristics measured were Total Sleep Time (TST), Total Sleep Episodes (TSE) and the Mean Sleep Time (MST) for each sleep episode.

Both *trbd^C518A^* and *w^1118^* females had less TST in LP (p<0.001) and more TST in DP (p<0.05) compared to males (Fig. 2C and Fig. 2D respectively). Females of *trbd^C518A^* and *w^1118^* were recorded to have significantly greater TSE in LP than their male counterparts (both p<0.0001) (Fig. 2E), while in DP differences were markedly reduced with *trbd^C518A^* females having a lower TSE than males (p<0.05) and *w^1118^* control males and females being statistically indistinguishable (Fig. 2F). Male flies showed more MST in LP compared to females (mutant p<0.001 and control p<0.0001) (Fig. 2G) but these differences were significantly reduced in DP. There, *trbd^C518A^* females showed more MST (p<0.05) than males while again, control males and females were statistically indistinguishable (Fig. 2H). These results suggested *de novo* sex dimorphism in DP for the *trbd^C518A^* mutants accompanied with fragmented sleep in mutant males (increase in TSE and decrease in MST) (Fig. 2F and Fig. 2H).

Focusing on the LP, introduction of the *trbd^C518A^* did not affect TST of mutant males (Fig. 2C) but increased their TSE (p<0.0001) (Fig. 2E). Within each TSE however, mutant male MST was significantly reduced compared to their wildtype counterparts (p<0.0001) (Fig. 2G). This indicated that mutant males had a fragmented sleep pattern in LP, with more sleep episodes but less sleep in each episode, when compared to the controls. In the females, introducing of the *trbd^C518A^* slightly increased TST and MST in LP (both p<0.01) (Fig 2C and Fig 2G) but no influence in TSE (Fig2E). In conclusion, the *trbd^C518A^* mutation caused more fragmented sleep patterns in males during the light period.

### Quantification of brain size and cells showed sex-specific changes in *trbd^C518A^* mutants

Our phenotypic data in this study suggested that there were neurodegeneration-associated symptoms in *trbd^C518A^* male flies. Our previous data indicated that deleting the *trbd* open reading frame (ORF) caused significantly elevated age-dependent vacuolisation in fly brains (Kounatidis et al., 2017). This was suppressed when NF-κB/Relish was silenced in glia (Kounatidis et al., 2017). To obtain histological evidence of potential early signs of cellular loss for *trbd^C518A^* mutants, we sought to quantify the neuron and glia content of whole brains through immunofluorescence. We dissected 18-day-old adult brains and stained them with DAPI (to visualise nuclei in all brain cells) and antibodies against Repo (labelling glia) or Elav (labelling neurons) (Fig. 3A). We then compared the projected brain size (Fig. 3B) and the signal levels of DAPI, Repo and Elav between wild type and mutant. Signal levels were measured as intensity and density. Intensity was the sum of fluorescent signal intensity from all Z stacks of single whole brains indicating the total number of cells (neurons, glia, or both) in the brain (Fig. C, E and G). The density of cells across the brain was the ratio of cell intensity divided by the projected brain size, indicating the distribution of the signal across the projected brain area (Fig. D, F and H).

**Figure 3.**
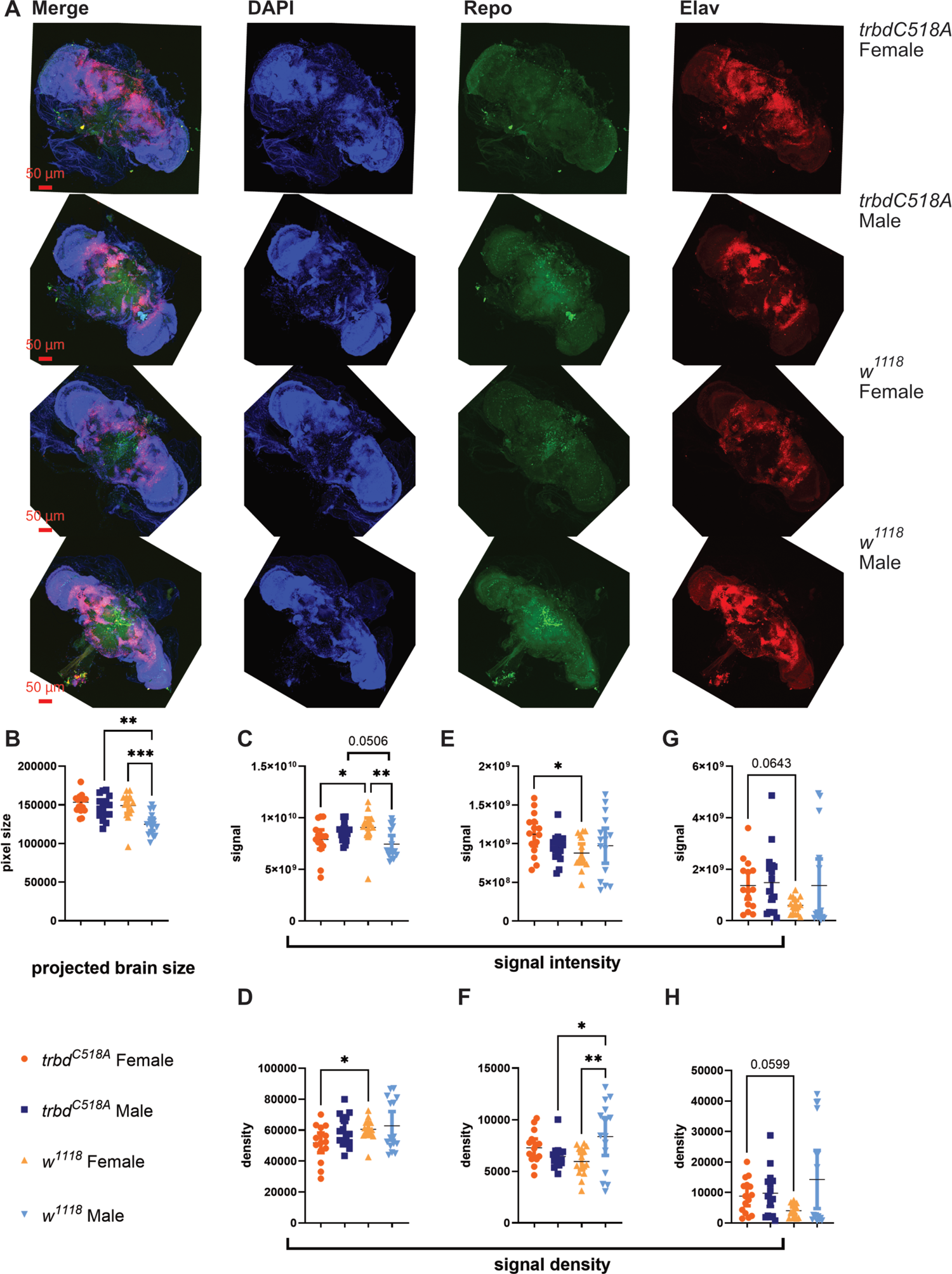
Quantification of neurons and glia by immunofluorescence indicate cell loss in *trbd^C518A^* males. **(A)** To quantify number of cells in 18-days old *Drosophila,* brains were stained using primary antibodies against Repo (for glia) and Elav (for neurons) and conjugated with corresponding secondary antibody. DAPI was used to counterstain the nuclei (scale bar 50 μm). **(B-H)** Sum of fluorescence signal intensity of every Z slice to measure projected brain area as well as fluorescence signal density of projected brain area (one-way ANOVA Kruskal-Wallis test with uncorrected Dunn’s test, n=15, 95% CI). **(B)** *w^1118^* males had a significantly smaller projected brain size compared to *w^1118^* females (***p<0.001) as well as *trbd^C518A^* males (**p<0.01). **(C)** *w^1118^* males had a significantly weaker DAPI signal intensity than *w^1118^* females (**p<0.01) and marginally weaker than *trbd^C518A^* males (p=0.0506). Female *trbd^C518A^* mutant flies had a significantly weaker DAPI signal intensity than *w^1118^* control females (*p<0.05) but were statistically indistinguishable from their male siblings. **(D)** *trbd^C518A^* females had a significantly weaker DAPI signal density than *w^1118^* females (*p<0.05). **(E)** *trbd^C518A^* females had a significantly stronger Repo signal intensity compared to *w^1118^* females (*p<0.05). **(F)** *trbd^C518A^* males had a significantly reduced Repo signal density compared to control *w^1118^* males (*p<0.05). This combined with the significantly higher projected brain size of the mutant males (see B, this figure) indicated loss of glia. Control *w^1118^* males had significantly higher Repo intensity from their female control siblings (**p<0.01). **(G)** *trbd^C518A^* females had marginally higher Elav signal intensity than *w^1118^* female (p=0.0643). **(H)** *trbd^C518A^* females had marginally higher Elav signal density than *w^1118^* female (p=0.0599).

*trbd^C518A^ has sex divergent impact in projected brain size, total cell quantity and densityw^1118^* males had a smaller projected brain area than *w^1118^* females (p<0.001) and both male *trbd^C518A^* mutants (p<0.01) (Fig. 3B). There was no significant difference in projected brain size between *trbd^C518A^* males and females (Fig. 3B), indicating that the sex bias of brain size in *w^1118^* controls was diminished when introducing the *trbd^C518A^* mutation. This conclusion was further supported by total cell quantification of DAPI signal intensity. *w^1118^* males had lower DAPI signal intensity than *w^1118^* female (p<0.01) and *trbd^C518A^* mutants (p=0.0506) (Fig. 3C). The smaller brain size of *w^1118^* male did not have a negative impact in its cell density (Fig. 3D). However, introducing *trbd^C518A^* in the females caused reduction in DAPI intensity as well as density compared to the same-sex control (both p<0.05) (Fig. 3C and Fig. 3D), without any significant difference in projected brain size (Fig. 3B). This suggested cell loss in the females after introducing the *trbd^C518A^* mutation.

*trbd^C518A^ positively affects female glia quantity and negatively impacts male glia densitytrbd^C518A^* females had stronger Repo intensity than control females (p<0.05) (Fig. 3E), indicating more glia cells in the mutant females, but there was no significance in Repo density (Fig. 3F). There was no significant difference when comparing Repo signal intensity in *w^1118^* males with *w^1118^* females and *trbd^C518A^* males (Fig. 3E). *w^1118^* males had elevated Repo signal density compared to *w^1118^* females (p<0.01) (Fig. 3F). *trbd^C518A^* males also had lower Repo signal density than their wild type counterparts (p<0.05) (Fig. 3F). These results suggested that *trbd^C518A^* positively influenced glia quantity in females while negatively impacting glia densities in males.

### trbd^C518A^ positively influences female neuron quantity and density

*trbd^C518A^* females had stronger Elav intensity (p=0.0643) (Fig. 3G) and density (p=0.0599) (Fig. 3H) compared to *w^1118^* females. This result indicated that the *trbd^C518A^* mutation could promote both neuronal as well as glia (see above) development in females. All other comparisons were statistically indistinguishable (Fig. 3G and Fig. 3H).

### Age-specific sex differences in brain immune activity are abolished when *trbd* is enzymatically inactivated

Deletion of the *trbd* ORF upregulated gene expression of Relish/NF-κB-dependent antimicrobial peptide genes (AMPs) (Fernando et al., 2014). Overexpression of AMPs in the brain caused neuroinflammation that was associated with NF-κB-mediated neurodegeneration (Cao et al., 2013; Kounatidis et al., 2017). Hence, the expression pattern of these AMP genes can potentially be used as hallmarks for analysing sex bias in neuroinflammation. Biochemically but also functionally, the *trbd* ORF deletion and the *trbd^C518A^* mutation are different. Deleting the *trbd* ORF will expose the ubiquitin chains to other DUBs that could then cleave them, while an enzymatically inactive Trabid will bind and protect the ubiquitin chains from other DUBs. This will eventually alter the Trabid-dependent NF-κB signal output without the potential involvement of other DUBs (see discussion). We used 3- and 18-day old adult fly brains to perform qPCR analysis on Relish-dependent AMP gene expression. The relative gene expressions of the control groups were used as baseline (baseline=1) for the relative fold changes of the test groups.

We did not record any differences in *trbd* expression across all samples (Figure 4A-F and Fig S1A-D). At day 18 of adulthood, *w^1118^* male brains had a lower AMP gene expression compared to *w^1118^* female brains. This was indicated by the relative expression of *attacin-A (attA)*, *attacin-B* (*attB*), *diptericin-A* (*dptA*), *cecropinC* (*cecC*) and *drosocin* (*dro*) (p<0.05) (Fig. 4A). This suggested sex-specific regulation but also age-dependent changes since at day 3, only *attB* was significantly lower (p<0.0001) in *w^1118^* male brain (Fig. S1A). However, there were no significant differences in AMP gene expression when comparing the mutant female and male brains at day 18 (Fig. 4A). At that time point, compared to sex-matched *w^1118^* controls, *trbd^C518A^* female mutants showed a reduced expression of *cecC*, *dro* and *dptA* (p<0.05) and *trbd^C518A^* males an elevated expression in *attA* (p<0.05), which levelled the two *trbd^C518A^* sexes (Fig. 4B). This result suggested that introducing the *trbd^C518A^* mutation eliminated the sexual dimorphism in AMP expression pattern observed in the genetic background. Nevertheless, the introduction of the *C518A* point mutation in *trbd* did cause significant upregulation of *attA* (p<0.05) and downregulation of *cecC* in *trbd^C518A^* females compared to controls of the same sex at day 3 (p<0.05) (Fig. S1B).

**Figure 4.**
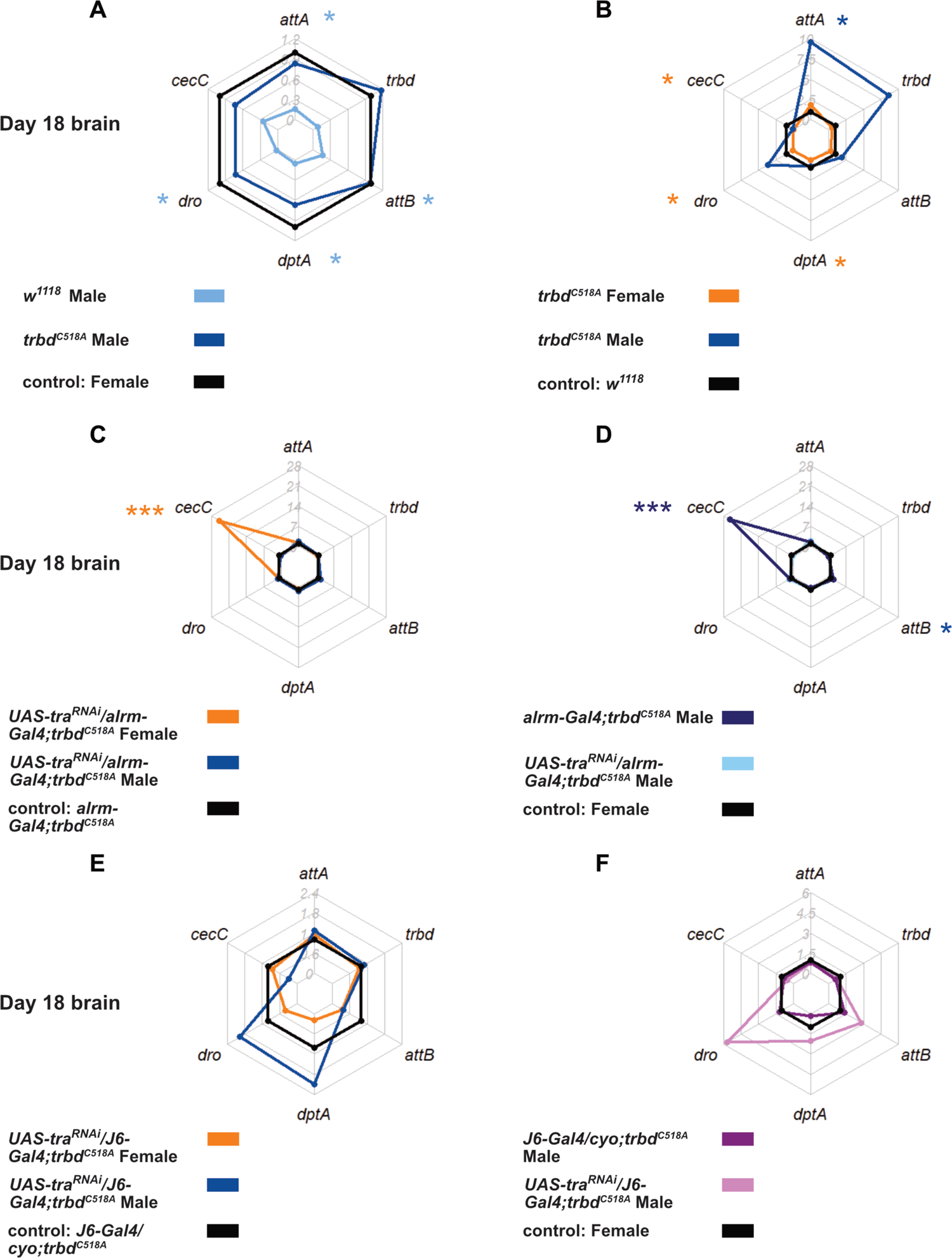
Sex-specific differences in AMP gene expressions are reduced when introducing the C518A mutation. **(A)** Normalized fold change of AMP gene expression in 18-days old *trbd^C518A^* and *w^1118^* male brains with *trbd^C518A^* and *w^1118^* females as the baseline control. **(B)** Normalized fold change of AMP gene expression in brains of 18-days old *trbd^C518A^* female and male flies with *w^1118^* females and males as the baseline control. **(C)** Normalized fold change of AMP gene expression in 18-days old *UAS-tra^RNAi^/alrm-Gal4; trbd^C518A^* female and male brains with *alrm-Gal4; trbd^C518A^* females and males as the baseline control. **(D)** Normalized fold change of AMP gene expression in 18-days old *UAS-tra^RNAi^/alrm-Gal4; trbd^C518A^* and *alrm-Gal4; trbd^C518A^* male brains with *UAS-tra^RNAi^/alrm-Gal4; trbd^C518A^* and *alrm-Gal4; trbd^C518A^* females as the baseline control. **(E)** Normalized fold change of AMP gene expression in 18-days old *UAS-tra^RNAi^/J6-Gal4; trbd^C518A^* female and male brains with *J6-Gal4/CyO; trbd^C518A^* females and males as the baseline control. **(F)** Normalized fold change of AMP gene expression in 18-days old *UAS-tra^RNAi^/J6-Gal4; trbd^C518A^* and *J6-Gal4/CyO; trbd^C518A^* male brains with *UAS-tra^RNAi^/J6-Gal4; trbd^C518A^* and *J6-Gal4/CyO; trbd^C518A^* females as the baseline control.

### Ubiquitylation and proteomic profiling suggests sex-specific effects of Trabid in neuronal and immune functions

Disruption in neuronal ubiquitin homeostasis has been involved in the pathogenesis of various neurodegeneration diseases (Hallengren et al., 2013). As detailed above, generating an enzymatically null *trbd* point mutant caused multiple sex-specific neurological phenotypes. Previous studies have provided insights in the involvement of Trabid in individual pathways, such as Imd/NF-κB (Fernando et al., 2014; Kounatidis et al., 2017), Wnt (Tran et al., 2008) and TLR (Jin et al., 2016) but the general picture remained unclear. We took advantage of rapid and deep-scale ubiquitylation and total proteome profiling to look at the global and sex specific ubiquitin landscape in 18-day old adult *Drosophila* heads (∼200 per sample with duplicates), in our model with wild type or enzymatically inactive Trabid groups (Steger et al., 2021)

We used ubiquitin GlyGly enrichment and liquid chromatography-mass spectrometry (LC-MS) to profile proteins that were enriched by ubiquitin modification and took 10% of lysate for total proteome analysis before the immunoprecipitation for housekeeping control (Steger et al., 2021). We identified significant differences when comparing sex-matched *trbd^C518A^* mutants to the *w^1118^* genetic background in GlyGly enrichment (Fig. S2A for females and Fig. S2B for males) as well as when comparing females to males within each genotype (Fig. S2C for *trbd^C518A^* and Fig. S2D for *w^1118^*). We also found differentially expressed proteins in total proteome when comparing sex-matched *trbd^C518A^* mutants to the *w^1118^* genetic background (Fig. S2E for females and Fig. S2F for males) as well as when comparing females to males within each genotype (Fig. S2G for *trbd^C518A^* and Fig. S2H for *w^1118^*).

Following these comparisons, results were subjected to Venn diagram analysis (Figure 5). We then searched each of the differentially expressed proteins (p<0.05 and fold change - log 2> or log2<) in the STRING proteome database to discover enriched or underrepresented pathways in relation to the *w^1118^* control. Comparing the *w^1118^* genetic background and the *trbd^C518A^* mutant within females and within males, identified pathways that were overrepresented (up) or underrepresented (down) in GlyGly enrichment (Fig. 5A and Table S1). Compared to *w^1118^,* flies carrying the *trbd^C518A^* mutation showed more differences in ubiquitylation of females (89 unique proteins) than males (38 unique proteins) (Fig. 5A and Table S1). Moreover, comparison between *trbd^C518A^* flies of different sex showed more ubiquitin modification differences (17 unique proteins) than between *w^1118^* females and males (10 unique proteins) (Fig. 5B and Table S1).

**Figure 5.**
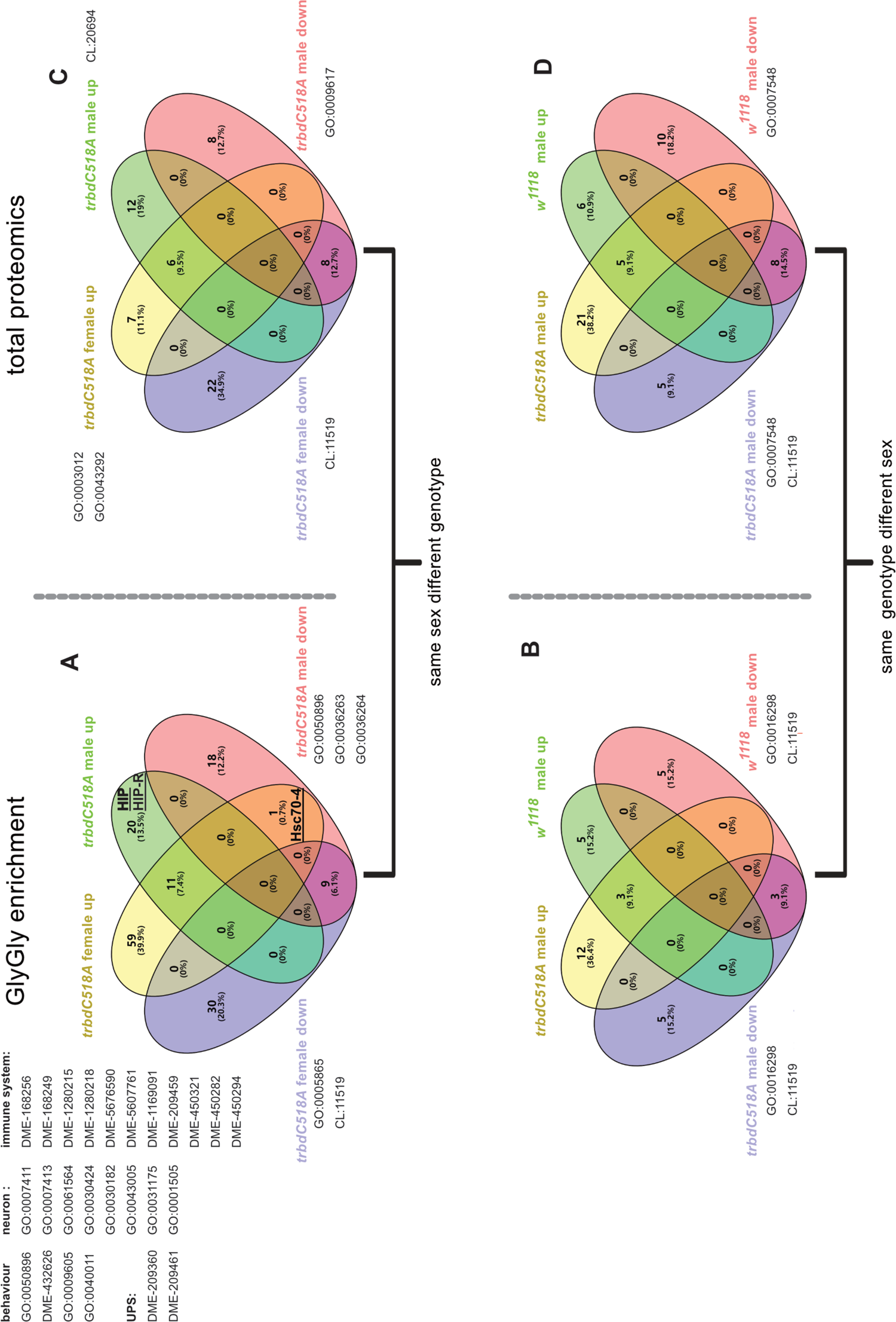
Ubiquitinomics and proteomics analysis reveals sex bias in gene ontology and interactome enrichment in the heads of *trbdC518A* and *w^1118^* flies. **(A)** Ubiquitin GlyGly enrichment Venn analysis by comparing differentially expressed protein hits (p<0.05 and log2<-1 or >1) between *trbd^C518A^* female or male to (respectively) *w^1118^* female or male. **(B)** Ubiquitin GlyGly enrichment Venn analysis by comparing differentially expressed protein hits (p<0.05 and log2<-1 or >1) between *trbd^C518A^* females and males or *w^1118^* females and males. **(C)** Total proteome Venn analysis by comparing differentially expressed protein hits (p<0.05 and log2<-1 or >1) when comparing between *trbd^C518A^* female or male and (respectively) *w^1118^* female or male. **(D)** Total proteome enrichment Venn analysis by comparing differentially expressed protein hits (p<0.05 and log2<-1 or >1) between *trbd^C518A^* females and males or *w^1118^* females and males.

At total proteome level, *trbd^C518A^* caused several differences in pathway enrichment (29 unique proteins for the females and 20 unique proteins for the males) when compared to *w^1118^* (Fig. 5C and Table S1). As with GlyGly, when comparing different sex in the same genotype for the total proteome, we found more differences in the mutants (26 unique proteins) than in *w^1118^* controls (16 unique proteins) (Fig. 5D and Table S1). In conclusion, the GlyGly assays indicated that female *trbd^C518A^* flies had a significantly increased level of ubiquitylation compared to their genetic background and became more divergent from their male counterparts. Some of these differences were reflected on the total proteome level (see below). These were relevant to neurodegenerative disease and neurological processes and are highlighted below.

Moreover, comparing proteins from ubiquitinome in proteome, we identified that the most significantly ubiquitylated proteins were not significantly affected, which indicated that ubiquitylation of these proteins was involved in regulatory processes (Fig. S4). There were a few significantly ubiquitylated proteins that were also significantly expressed, mainly yolk (Yp) family proteins that are related to female sex development (Fig. S4 A, C and D). This suggested that the ubiquitylation of Ypproteins were related to expression. Finally, there were proteins only significantly expressed but not significantly ubiquitylated, which implied that ubiquitylation was mediating stabilisation or degradation of these proteins (Fig. S4).

### Hsc70-4 is differentially regulated in females and males

Hsc70-4 is a protein involved in protein folding and has been suggested to play a role in several neurodegenerative disease models in *Drosophila* (Iijima-Ando *et al.,* 2005; Johnson *et al.,* 2020). Moreover, in mice it was found to coordinate the homeostasis of microtubule and tau accumulation (Fontaine *et al.,* 2015). In human cells, the E3 ligase CHIP-Hsc70 complex and UbcH5B selectively modify phosphorylated tau (Shimura et al., 2004). Our data revealed that Hsc70-4 regulation by ubiquitin modification was significantly divergent between females and males. Relative to *w^1118^* and compared to sex-matched controls, Hsc70-4 was downregulated in *trbd^C518A^* males while upregulated in *trbd^C518A^* females (underlined in Fig. 5A). Moreover, two H c70-interacting proteins (HIP and HIP-R) located in the X chromosome were exclusively upregulated in ubiquitylation when comparing *trbd^C518A^* mutant males to *w^1118^* males (underlined in Fig 5A). This implied that a dysfunction in an autosomal DUB, Trabid, can cause potentially direct or indirect divergent ubiquitylation on proteins coded in the X-chromosome.

### Sex bias in pathways related to locomotor activity

We found “muscle component” (GO:0005865) was less ubiquitylated in the female mutants than female controls (Fig. 5A and TableS1) and we detected enrichment of “muscle function” (GO:0003012 and GO:0043292) in the total proteome list for the female mutants (Fig. 5B and TableS1). This suggested that changes in ubiquitylation of muscle components and consequently enhanced muscle function might help female mutants to retain the same level of locomotor activity (Fig. 2B). When inactivating Trabid, proteins related to “response to stimulus” (GO:0050896) were less ubiquitylated in the male mutants and enriched in the female mutants (Fig. 5A and Table S1). This result could explain why C518A mutation had more impact on male locomotion (Fig. 2B).

### Female specific ubiquitylation in circadian rhythm

Inactivated Trabid caused more fragmented sleep pattern in males than in females (Fig. 2E-H). When compared to *w^1118^* females, reactome term “circadian clock pathway” (DME-432626) was significantly enriched in *trbd^C518A^* females (Fig. 5A and Table S1), suggesting when Trabid was inactivated, ubiquitylation in female “circadian clock pathway” may ameliorate negative influence on sleep pattern.

### Sex dimorphism in neuron development and neurotransmitter presence

Our histology results indicated sex-specific changes in cerebral cell quantity and density (Fig. 3). When introducing *trbd^C518A^* in female flies, we recorded GOs related to axon (GO:0007411, GO:0007413, GO:0061564 and GO:0030424) and neuron (GO:0030182, GO:0043005 and GO:0031175) were enriched by ubiquitin modification (Fig. 5A and Table S1). This could explain our observation of increased glia and neuron quantities and densities in mutant females. We noticed the GO “regulation of neurotransmitter levels” (GO:0001505) (Fig. 5A and Table S1) was enriched by ubiquitylation. GOs related to dopamine monooxygenase (DBH) activity (GO:0036263 and GO:0036264) were less ubiquitylated in *trbd^C518A^* males compared to *w^1118^* males (Fig. 5A and Table S1). DBH is involved in the synthesis of the neurotransmitter norepinephrine. Sex specificity in ubiquitin modification of neurotransmitters could explain the sex bias in neurological behaviours.

### Divergence in immunity between males and females

We detected sex-based divergence in Relish-induced AMP gene expression in the brain of mutants when compared to *w^1118^* controls but also compared to each other (Fig. 4A and Fig. 4B). In contrast, there was no sex-specific enrichment at the level of the pathway (Fig. 5B and Fig. 5D). Nevertheless, introducing the *trbd^C518A^* mutation in female flies, produced more reactome hits related to immunity (DME-168256, DME-168249, DME-1280215 and DME-1280218), including NF-κB signalling (DME-5676590, DME-5607761 and DME-1169091) Imd (DME-209459) and JNK-MAPK (DME-450321, DME-450282 and DME-450294). These were recorded as significantly enhanced by ubiquitylation (Fig. 5A and Table S1). Given the fact that ubiquitylation is heavily involved in immune-related pathways, we were not surprised to find ubiquitylation-related proteins were dominant components in these hits (Table S1). At the total proteome level, we recorded that the hit “response to bacterium” (GO:0009617) was downregulated (Fig. 5C and Table S1) and AMP — AttA or AttB was upregulated (Table S1) in mutant males. Elevation of *attA* gene expression was recorded in male mutant brains by qPCR (Fig. 4B). The *Drosophila* Sting pathway has been reported to co-regulate Relish-induced AMP gene expression with the canonical Imd pathway (Martin et al., 2018). Three Tot family protein as well as Sting were matched to GO:0009617 (Fig. 5C, Table S1). In total, ubiquitylation had a larger influence in immune regulation when *trbd^C518A^* was introduced in female flies than in the male flies, but the output of total proteome suggested that the immune response was altered in males but not in females.

### Sex dimorphism in ubiquitin linkages and ubiquitin-proteasome system (UPS)

Because Trabid is a Lys29 and 33 linkage-specific DUB (Fig. 1C), we investigated the pathway changes in the ubiquitin system. We recorded enrichment of ubiquitin-proteosome system (UPS) associated reactome hits, including DME-209360 (ubiquitination and proteolysis of phosphorylated CI) and DME-209461 (ubiquitination and degradation of phosphorylated ARM), in female mutants, compared to sex-matched *w^1118^* controls (Fig. 5A and Table S1). Polyubiquitin (Ubi-p63E), ubiquitin thioesterases (CG4603 and CG4968), E2 enzyme (eff) and component of proteasome (Rpn8 and Prosbeta3) (Table S1) were enriched UPS-related proteins in the mutant females, when compared to *w^1118^* females. This indicated that inactivated Trabid could elevate the UPS activity in female heads. When introducing *trbd^C518A^*, we did not record any pathway hits associated to ubiquitination (Fig. 5A) in males, but we recorded polyubiquitin (Ubi-p63E) accumulation (Table S1). Taken together, these findings strongly suggest that sex-specific ubiquitylation took place under an inactive Trabid.

Except M1 and Lys29 linkages, the GlyGly ubiquitin enrichment allowed us to relatively compare ubiquitin linkages between two samples. We found Lys6, 27 and 33 linkages were accumulated in the mutant female heads (p<0.05, Fig. 6A-C). *In vitro,* Lys29 and 33 chains were the primary substrates for Trabid hydrolysis (Fig. 1B). This accumulation was not observed in the mutant males. The latter result suggested that the C518A mutation had a sex-specific impact in Lys6, 27 and 33 homeostasis.

**Figure 6.**
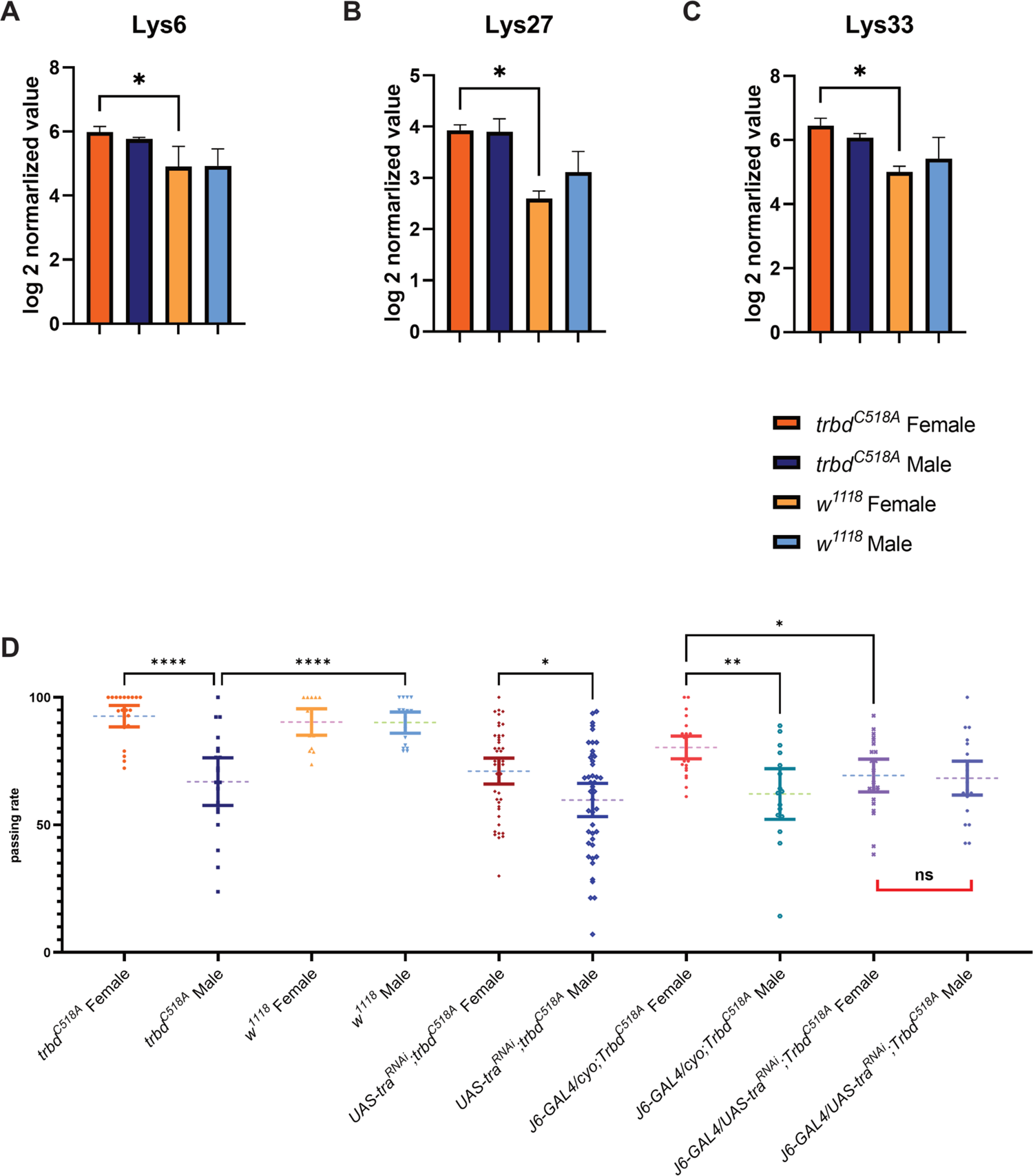
*trbd^C518A^* influences Lys6, 27 and 33 homeostasis while blocking female sex development in *trbd^C518A^* astrocytes and immunocompetent cells reduces sex bias in climbing. **(A)** Heads of *trbd^C518A^* females had higher levels of Lys6 ubiquitin than *w^1118^* females (one-way ANOVA Kruskal-Wallis test with uncorrected Dunn’s test, n=2, SD). **(B)** Heads of *trbd^C518A^* females had higher levels of Lys27 ubiquitin than *w^1118^* females (one-way ANOVA Kruskal-Wallis test with uncorrected Dunn’s test, n=2, SD). **(C)** Heads of *trbd^C518A^* females had higher levels of Lys33 ubiquitin than *w^1118^* females (one-way ANOVA Kruskal-Wallis test with uncorrected Dunn’s test, n=2, SD). **(D)** Climbing assay of *trbd^C518A^, w^1181^, UAS-tra^RNAi^; trbd^C518A^, J6-GAL4/CyO; trbd^C518A^* and *J6-GAL4/UAS-tra^RNAi^; trbd^C518A^* (one-way ANOVA Kruskal-Wallis test with uncorrected Dunn’s test, n=approx. 300 flies, 95% CI).

### Cell-specific blocking of female development reduced sex bias in neurological and inflammation phenotypes of the *trbd^C518A^* mutation

The gene *transformer* (*tra*) is a member of the female somatic sexual differentiation pathway (Baker and Ridge, 1980). Mutations in *tra* transform chromosomal females (X/X; *tra/tra*) into sterile males (Butler et al., 1986). This is because a functional Tra protein is produced in females, but not males (Sosnowski et al., 1989). Our data so far indicated, that *trbd^C518A^* flies had sex-specific neurodegeneration-related traits as we were able to detect early locomotion (climbing) defects (day 6) and loss of glial density (day 18) in *trbd^C518A^* males (Fig. 2B and Fig 3F). Moreover, we detected upregulation of male AMP gene expression related to controls (Fig 4A). Using the Gal4-UAS system we reduced, through RNAi, *trbd^C518A^* female sex development immunocompetent cells (fat body, gut, glia, and haemocytes) (*J6-GAL4,* Glittenberg et al., 2022) and carried out the climbing assay 9 days after eclosion to explore if the sex bias would be reduced. We used *J6-GAL4/CyO; trbdC^518A^* as controls.

As expected, we observed a slower climbing in control *trbd^C518A^* males compared to their female siblings (Fig. 6D). When *tra* was silenced in female immunocompetent cells, sex-dependent differences in climbing ability were diminished (Fig. 6D). Specifically, when suppressing female development in *trbd^C518A^* immunocompetent tissues, female climbing ability decreased to levels statistically indistinguishable from the males (Fig 6D).

Since the *Drosophila* IMD pathway has disparate regulation in different sexes and tissues (Vincent and Dionne, 2021), we studied the expression of AMP genes in a *trbd^C518A^* background where female development was reduced through *tra* RNAi. This time we used an additional GAL4 line that drives expression in a sub-population of glia cells (astrocytes) (*alrm-GAL4*; Weiss *et al,* 2022). We used *J6-GAL4/CyO; trbd^C518A^* and *alrm-GAL4/CyO; trbd^C518A^* as controls. When reducing female development in astrocytes (*alrm-GAL4/UAS-tra^RNAi^; trbd^C518A^*), *cecC* expression levels were elevated in the 18-days old female mutant brain compared to the female control (Fig. 4C). We further investigated the difference between males and females. In the controls (*alrm-GAL4; trbd^C518A^*), males had significantly higher *cecC* (p<0.001) and lower *dptA* (p<0.05) expression compared to their female siblings (Fig. 4D). In *alrm-GAL4/UAS-tra^RNAi^; trbd^C518A^*, these sex-specific differences in *cecC* and *dptA* were recorded at day 18 (Fig. 4D) but not at day 3 (Fig. S1C and Fig. S1D). The finding suggested that supressing female development in astrocytes reduced the sexually dimorphic expression of AMPs in the brain in an age-dependent manner.

We also measured the consequences of silencing *tra* on brain AMP gene expression in *J6-GAL4/CyO; UAS-tra^RNAi^; trbd^C518A^*. Expressing *tra^RNAi^* in immunocompetent cells of *trbd^C518A^* through the *J6-GAL4,* caused no significant differences in AMP gene expression in any comparison in the brain (Fig. 4E and Fig. 4F). This indicated that supressing female development did not have an impact on AMP expression by silencing *tra* through the *J6-GAL4*.

## Discussion

It has been established that females and males have different vulnerabilities to neurodegenerative diseases like AD and PD (Palta et al., 2021; Rajan et al., 2021). However, the molecular mechanisms that contribute to this sex-specificity remain largely unexplored. Recently, the X-linked ubiquitin specific peptidase 11 (USP11) was found to augment pathological tau aggregation via tau deubiquitination initiated at Lysine-281 in female mice and AD human brain tissue (Yan et al., 2022). This, as well as previous studies, brought forward the idea that the balance of ubiquitinylation leading to protein degradation may be disrupted in AD and PD leading to protein accumulation. At the same time, Genome Wide Association Studies (GWAS) implicate numerous innate immune genes as risk factors for late onset AD in a glial transcriptional network (Kosoy et al., 2022; Matarin et al., 2015; Salih et al., 2019; Sims et al., 2017). This suggests that genetic differences (as defined by GWAS) in the response to the normal age-dependent increase in inflammatory activity may lead to late onset AD (Fulop et al., 2021 for review). Central to this age-dependent inflammatory activity is NF-κB signalling (Kaltschmidt et al., 2022 for review). Here, using *Drosophila*, we align these three aspects of neurodegeneration: the sex-specific variation, the balance of ubiquitylation, and the role of NF-κB-regulated inflammation.

Trabid is a member of the A20 family DUBs, which include A20, Trabid and Cezanne (Evans et al., 2001). In mammals, A20 is a negative regulator of IKK/NF-κB activation (Lork et al., 2017). In *Drosophila,* knocking out the *trbd* ORF leads to over activation of IKK-induced NF-κB signalling (Fernando et al., 2014; Hua et al., 2022; Kounatidis et al., 2017). A20 mediates Met1, Lys48 and Lys63 ubiquitin chains to regulate inflammation (Lork et al., 2017; Wertz et al., 2015) and Trabid can regulate Lys63 ubiquitin chains *in vitro* (Fig 1D) and *in vivo* (Fernando et al., 2014; Licchesi et al., 2012). It is tempting to hypothesize that *Drosophila* Trabid is substituting A20’s role in NF-κB signalling since we know that its loss triggers glia-dependent inflammation, which induces neurodegeneration (Kounatidis et al., 2017) *In vitro*, DUB assays showed that hTrabid specifically hydrolyses Lys29 and 33 chains (Licchesi et al., 2012), which are rarely studied *in vivo*, and less active towards Lys63 chains. Our DUB assays showed that dTrabid had the same enzymatic activity as hTrabid. Moreover, *trbd^C518A^* mutation led to Lys33 ubiquitin accumulation in female heads, indicating the unique role of Trabid in regulating ubiquitin homeostasis *in vivo*.

*In vivo*, the enzymatic inactivation of Trabid, resulted in larger male brains when measured in 18-day old flies (Fig. 3B). Despite that, glial density in the *trbd^C518A^* male brains was reduced compared to the male controls (Fig. 3F). This indicated mutant male brain cell loss and correlated with the neurological phenotypes of *trbd^C518A^* males where reduction of climbing ability was observed early in 9-day old flies and deteriorated at day 18 (Fig. 2B). Moreover, significantly increased fragmented sleep was recorded at day old 7 for *trbd^C518A^* males (Fig. 2C). In contrast, we recorded a gain of glia in the *trbd^C518A^* female brains since they had a significant increase in signal intensity compared to their female controls (Fig. 3E). In addition, female brains showed a marginal gain of neurons (Fig. 3G and Fig. 3H) and an enrichment in neuron-related GO terms in mutant female brains compared to their sex-matched controls (Fig. 5A). This correlated with the fact that the mutant females had a significantly higher climbing ability than their male siblings while they were statistically indistinguishable from the sex-matched controls (Fig 2B). These neurological phenotypes were underscored by the increase of AMP gene expression in *trbd^C518A^* males and the decrease of AMP gene expression in *trbd^C518A^* females (Fig. 4A and Fig. 4B). Noteworthy, the level of ubiquitylated Hsc70-4, which has a protective role in neurodegeneration (Shimura et al., 2004), was upregulated in mutant females and downregulated in mutant males, when comparing to controls (Fig 5A). While two Hsc70-interacting proteins’ ubiquitylation levels were upregulated in mutant males, when compared to control males (Fig 5A). Moreover, inactivating *trbd* only caused accumulation of Lys6, 27 and 33 ubiquitin linkages in females (Fig 6A-C). Ubiquitylation profiling suggested that the sex divergence in neurological phenotypes of *trbd^C518A^* is associated to differences in ubiquitylation of X-chromosome proteins and ubiquitin linkage homeostasis. As an autosomal DUB, loss of Trabid enzymatic activity results in sex-specific neurodegeneration and early neurological phenotypes while changing the inflammatory profile of the two sexes. Tissue-specific silencing of female somatic sexual differentiation pathway in *trbd^C518A^* via *tra^RNAi^* revealed sex-specific difference in climbing defects (Fig 6D) and brain inflammatory can be reduced (Fig 4C and Fig 4D). More work is needed to identify the mechanistic connection of Trabid with neuronal pathways identified by GlyGly/LCMS that connect loss of brain function to sex-specific neurodegenerative phenotypes.

## Material and Methods

### Alignment

The amino acid sequence of *Homo sapiens, Mus musculus* and *Drosophila melanogaster* were obtained from UniProt (https://www.uniprot.org/). The multiple sequence alignment was done by Clustal Omega (https://www.ebi.ac.uk/Tools/msa/clustalo/) and visualize by Jalview 2.11. The hTrabid-AnkOTU’s crystal structure 3ZRH and dTrabid-AnkOTU’s AlphaFold2 predicted structure (319-751) were imported and aligned in PyMol.

### Plasmid cloning

For cloning dTrabid-AnkOTU into poPINS plasmid, Primer “GCGAACAGATCGGTGGTGACACCCTGCAAGAGCGTCAAGAGAG” and “ATGGTCTAGAAAGCTTTATTCTTCGTCGGAGTCGCCGTCGG” were used to amplify dTrabid-AnkOTU domain. We synthesized the dTrabid gene with codon optimization to *Spodoptera frugiperda* (Sigma, UK Suppl) and cloned into HindIII and KpnI digested poPINS plasmid. Positive colonies were selected by beta-gal and confirmed by Sanger sequencing (Sourcebioscience, UK). We used “TTCTGCTGGCGACGCCCTGCTGGACTCTGC” and “TGCATAGCAGAGTCCAGCAGGGCGTCGCCA” to create the C518A mutation with the original cloning primers. The sequence was confirmed by Sanger sequencing (Sourcebioscience, UK).

### Protein purification

Plasmid was transformed into Rosetta II E. coli competent cells and selected on kanamycin and chloraphenicol LB plate. Respectively, all colonies from on the LB plate were made into 300 ml overnight LB liquid preculture which contains 30 µg/ml kanamycin and 34 µg/ml chloraphenicol. Every liter of 2xTY media were inoculated with 15 ml overnight preculture with 30 µg/ml kanamycin and 34 mg/ml chloraphenicol. The cultures were incubated at 30 for 3 hours and temperature was lowered to 18 and 400 mM of IPTG were added and 100 mM ZnSO4 was added to Trbd-NZF1 and -NZF cultures. The cultures were incubated overnight, harvested, and stored in −80.

Harvested cell pellets were defrosted at room temperature (23) and sonicated for 2 minutes at 30 amp (15 seconds impulse and 15 seconds rest) in lysis buffer (20 mM tris, 300 mM, 20 mM imidazole, 2 mM ethanethiol and pH 8). Lysate was centrifuged for 30 minutes at 15000 rpm at 4. Lysate was subjected to Ni-column purification and eluted with 20 mM tris, 300 mM, 500 mM imidazole and 2 mM ethanethiol (pH 8). Elution was dialysed with anion wash buffer (20 mM Tris (pH 7.5) and 4 mM DTT) with 1 mg SUMO protease at 4 for overnight.

Ni-column purified elutes were subjected to ResourceQ anion exchange chromatography. The wash buffer contained 20 mM Tris (pH 7.5) and 4 mM DTT and elution buffer contained 20 mM Tris (Ph 7.5), 4 mM DTT and 1 M NaCl. Elution fractions were analysed by SDS-PHAGE gel. Fragments that were about 48 kDa were collected and concentrated.

Concentrated fragments were subject to SuperDex75 gel filtration chromatography. The gel filtration buffer contained 20 mM Tris (pH 7.5), 4 mM DTT and 750 NaCl. Filtered fragments were analysed by SDS-PAGE gel. Fragments that contained solo band that were 48 kDa were pooled, concentrated and snap freeze for long term storage.

### Deubiquitinase assay

Deubiquitinase was diluted to 300 or 600 nM using gel filtration buffer (20 mM Tris, 4 mM DTT and 750 NaCl). 10x DUB buffer that contained 500 mM, 500 mM NaCl and 50 mM fresh DTT. 2x master mix that 18μl water, 6μl 10 x DUB buffer and 6μl diubiquitin chains (0.6μg/ml) were prepared for each ubiquitin chain type and time point and the finale concentration of diubiquitin chains were 600 nM. We used 300 nM dTrabid-AnkOTU to hydrolysis all eight diubiquitin chains for 0, 5 and 30 minutes. We used 600 nM dTrabid-AnkOTUC518A to hydrolysis Lys29 and 63 diubiquitin chains for 0, 5 and 30 minutes. The result was visualized using protein gel electrophoresis and silver staining (Bio-rad, UK).

### Drosophila stock and crossing

Rainbow Transgenic Fly Inc (CA, USA) generated via CRISPR-Cas9 the *trbd^518C2A^* flies in a *w^1118^* genetic background, and we received 4 individual lines. For each line, males were balanced by crossing with *TM3^Sb^/Dr*, F1 virgins and F1 males that were *TM3^Sb^* were crossed and the F2 progeny were used for establishing stocks. From these we selected progeny that had no *TM3^Sb^* marker and verified the point mutation using primers “accgcgacgcgcagaaaca” and “ctcgtactctttccagcgag”.

To block female development in a tissue-specific manner we used *UAS-tra^RNAi^* (VDRC#2560). *Gla/CyO;trbd^C518A^/TM6B* was used to cross with *UAS-tra^RNAi^/CyO;TM6B/pri*, *and then with alrm-Gal4/CyO;TM6B/pri* or *J6-Gal4/CyO;TM6B/pri* to generate *UAS-tra^RNAi^/alrm-Gal4;TM6B/trbd^C518A^* or *UAS-tra^RNAi^/J6-Gal4; TM6B/trbd^C518A^*. Flies were collected at the day of eclosion and 20 flies were kept in a vial. Food was changed every three days.

### Climbing assay

*trbd^C518A^* and *w^1118^* were used for the assay. The assay was set up according to a recent protocol (Madabattula et al., 2015). Flies needed to climb 17.5 cm in a transparent graduate cylinder in 2 minutes to pass the assay. Around 20 flies were tested each time and the passing ratio was calculated by the equation:

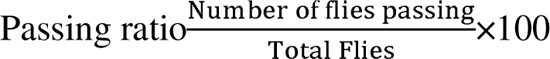

Flies were reared in 25 and 34% humidity. Assay were performed at room temperature (23) and in the afternoon (1 pm – 5 pm). In GraphPad Prism 9.2.0, normal distribution was checked for each group and two-way ANOVA was performed, using mixed-effects model with Tukey’s multiple comparison correction.

### *Drosophila* activity monitor (DAM)

On day-5 of adulthood, flies were anesthetized on ice and loaded on DAM with fresh food. The incubators were set at 25. The recording starts on day 6, with the light period (LP) starting from 8 am and dark period (DP) starting from 8 pm. The recording lasted for at least 3 days. We chose the second day (7 days after eclosion) data for analysis. In GraphPad Prism 9.2.0, normal distribution was checked for each group. One-way ANOVA Kruskal-Wallis with uncorrected Dunn’s test was applied for others. Multiple comparisons were conducted and plotted out.

### *Drosophila* lifespan assay

∼20 flies were kept in a vial and food was changed every three days and dead flies were counted. Food change lasted until the last fly’s death in the vial. In GraphPad Prism 9.2.0, we used built-in survival analysis to perform Mantel-Cox (Log-rank) test, calculate the median survival and visualize the result.

### RNA isolation and cDNA synthesis

*Drosophila* were reared to day 3 and 18 after eclosion, by the same method we used for lifespan. Brains were dissected in PBS buffer and RNA was isolated immediately using TRIzol Plus RNA Purification System (ThermoFisher, UK). Samples were stored in −80. We used Superscript III cDNA synthesis kit (ThermoFisher, UK) to synthesize cDNA.

### qPCR

We used SensiFast (Bioline, UK) to carry out the qPCR reaction. Gene *sdha* was the house keeping gene. The primers were listed in table below. We used CFX Connect Real-Time PCR system (Bio-rad) to collect ct values. 45 amplification cycles were conducted for each run. We used the R package “qpcr” (https://cran.r-project.org/web/packages/qpcR/qpcR.pdf) to perform “t-test” analysis and “relative expression” following the user manual. We used the R package “fmsb” to present the relative expression in spider plots.

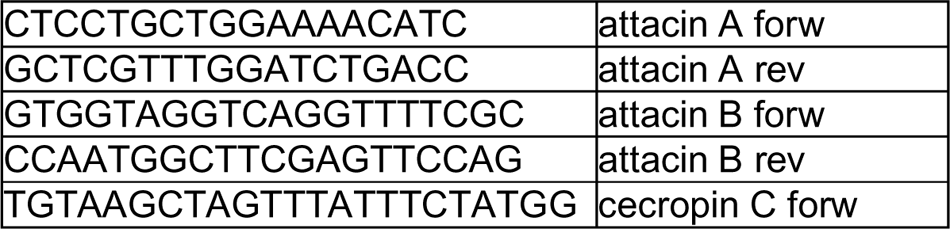
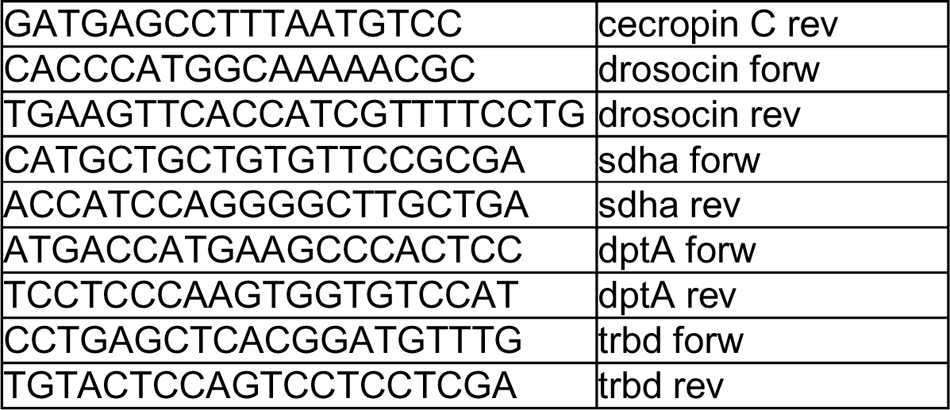

### Brain dissection and slide preparation

Three biological replications were prepared in total, and each replication had at least 5 brains. Flies were anesthetized on ice at day-18 after eclosion and dissected in PBS. PBTX solution (PBS and 0.3% Tween 80) and 4% PFA were used to prepare fix solution. Brains were moved to fix solution and fixed at room temperature for 30 minutes incubation. Samples were washed for 3 times in PBXT and transferred to Intercept (PBS) blocking buffer (Li-Cor, UK) for an hour incubation at 4 degrees with gentle rocking. Primary antibodies (1%, Rat-Elav-7E8A10 anti-elav and 8D12 mouse anti-Repo, DSHB, USA) in PBTX was used to replace the blocking solution and incubated at 4 degrees for overnight with gentle rocking. On the second day, samples were washed with PBTX for three times and replaced with secondary antibody (1%, goat anti-mouse Alexa Fluor 488 and goat anti-rat Alexa Fluor 568, ThermoFisher UK) and 1 ug/ml DAPI solution. After 45 minutes, samples were washed in PBTX solution for 3 times and incubated in PBXT solution for more than 45 minutes. Samples were incubated in Vectashield mounting media (2BScientific, UK) for more than 30 minutes and mounted on glass slide with cover slip.

Specimens were imaged on an Olympus SpinSR10 spinning disk confocal system equipped with Prime BSI sCMOS camera. 20x dry objective (0.8NA, UPLXAPO20X) was used to acquire the images. The microscope was driven using Olympus cellSens Dimension software. The image data were uploaded and stored in the Department of Biochemistry, University of Oxford OMERO server.

### Histology analysis

Images were analysed in FIJI-ImageJ. We summed fluorescence intensity across Z slices to quantify immunofluorescence signal. Raw images were background subtracted with the rolling ball subtraction method in ImageJ prior to the intensity calculation. Brain areas were calculated by outlining the DAPI channel signals. In each channel, the area and raw integrated density (RID) of the brain and background were measured. The signal intensity of the brain (foreground) of each channel was calculated by the equation:

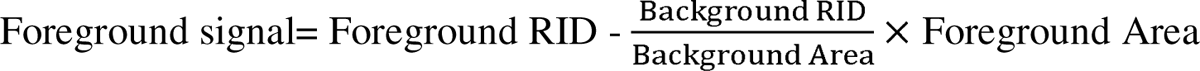

The foreground signal density of the brain was calculated by the equation:

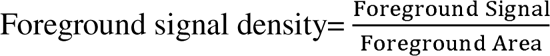

We used GraphPad Prism built-in normality test to test the distribution. One-way ANOVA and Kruskal-Wallis with uncorrected Dunn’s test was applied for others. Multiple comparisons were conducted and plotted out.

### Fly protein isolation

*Drosophila* were collected at the day of eclosion. 20 flies were kept in a vial and food was changed every 3 days. ∼200 flies were flashed froze by dry ice on day-18. Fly heads were cut off and placed into lysis buffer from PTMScanHS Ubiquitin/SUMO Remnant Motif Kit Protocol (#59322, Cell Signaling, UK), that contained 5% SDS in TEAB buffer (T7408, SigmaAldrich, Germany). Samples were homogenized by an electric motor pestle for 2 minutes at room temperature, sonicated following protocol instruction and centrifuged for 15 minutes at 12000 rpm. The supernatant was transferred into a new 1.5 ml microcentrifuge. The concentration of protein was measured by Pierce 660 assay (ThermoFisher, UK) with Ionic Detergent Compatibility Reagent (ThermoFisher, UK). Protein samples were flashed froze for further GlyGly ubiquitin enrichment experiment.

### GlyGly ubiquitin enrichment

Profiling of protein ubiquitylation by anti-GlyGly enrichment was performed as described before (Pinto-Fernandez et al., 2021)). In brief, we normalized all samples’ input to 1 mg. We followed the protocol of PTMScan HS Ubiquitin/SUMO Remnant Motif Kit Protocol (#59322, Cell Signalling, USA) to perform the Reduction and alkylation of protein and S-trap clean-up protease digestion. After trypsin digestion, 50μl (∼30μg) of aliquot was taken from the resuspension and used for total protein LCMS analysis. We continued to perform immunoaffinity purification (IAP) following the PTMScan HS Ubiquitin/SUMO Remnant Motif Kit Protocol.

### LC-MS analysis

Tryptic peptides were analysed using a Dionex Ultimate 3000 nano UPLC (Thermo Scientific) coupled to an Orbitrap Fusion Lumos Tribid mass spectrometer (Thermo Scientific). Briefly, peptides were trap on a PepMap C18 trap columns (Thermo) and separated on an EasySpray column (50cm, P/N ES803, Thermo) over a 60-minute linear gradient from 2 % buffer B to 35 % buffer B (A: 5 % DMSO, 0.1 % formic acid in water. B: 5 % DMSO, 0.1 % formic acid in acetonitrile) at a flow rate of 250 nL/min. The instrument was operated in data-independent acquisition mode as previously described in Steger M. et al. 2021 (Steger et al., 2021).

Proteins and GG-containing pepdtides were identified using the DIA-NN search engine (version 1.8), using the Uniprot database for Drosophila melanogaster (Taxon ID. 7227) and data visualisation was performed in Perseus (version 1.6.2.3). Volcano plots were generated using protein group intensities and GG-peptide intensities (precursor ions were collapsed into peptides following instructions from Steger M. et al. 2021 (Steger et al., 2021)) from DIA-NN.

We mapped 1388 ubiquitinated proteins. We kept the reads that had valid values in both biological replicates and resulted in 944 proteins. Quality control suggested the processed data set was suitable for further statistical analysis. Two sample t-test was performed, and we used proteins that were FDR>0.05 and log fold change<-1 or >-1 to perform Venn diagram analysis. The Venn diagram analysis was performed in Venny 2.1 (https://bioinfogp.cnb.csic.es/tools/venny/). Results were further analysed using the STRING databank (https://string-db.org). We did a Venn diagram analysis on glygly enriched proteins and total proteins. We selected the 1550 proteins that were exclusive to total proteins and adding the ubiquitin linkages result for width adjustment normalization analysis.We further matched the ubiquitylated protein with expressed proteins (fig. S4). The match ratios were about 97%. The mass spectrometry proteomics data will be made available via the ProteomeXchange Consortium via the PRIDE partner repository (Perez-Riverol, Y et al., 2019)

## Acknowledgements

We thank the Discovery Proteomics Facility led by Roman Fischer and Iolanda Vendrell for expert help with mass spectrometry analysis. APF and BMK were supported by the Chinese Academy of Medical Sciences (CAMS) Innovation Fund for Medical Science (CIFMS), China (grant number: 2018-I2M-2-002) and by Pfizer. ID was supported by Wellcome, PE was supported by an MRC Career Development Grant and PL was supported by the Royal Society (IEC\NSFC\201298) and the John Fell Fund (0010611).

---

**Figure S1. Differences in AMP gene expression (A)** Normalized fold change of AMP gene expression in 3-day old *trbd^C518A^* and *w^1118^* male brains with *trbd^C518A^* and *w^1118^* females (respectively) as the baseline control. **(B)** Normalized fold change of AMP gene expressions in 3-day old *trbd^C518A^* female and male brains compared to *w^1118^* female and male brains (respectively) as the baseline control. **(C)** Normalized fold change of AMP gene expression in 3-day old *UAS-tra^RNAi^/alrm-Gal4; trbd^C518A^* female and male brains with *alrm-Gal4; trbdC518A* female and male (respectively) as the baseline control. **(D)** Normalized fold change of AMP gene expression in 3-day old *UAS-tra^RNAi^/alrm-Gal4; trbd^C518A^* female and *alrm-Gal4; trbd^C518A^* female brains with *UAS-tra^RNAi^/alrm-Gal4; trbd^C518A^* male and *alrm-Gal4; trbd^C518A^* male (respectively) as the baseline control.

**Figure S2. Volcano plots of ubiquitin GlyGly and total proteome comparisons. (A)** Significant protein hits from ubiquitin GlyGly enrichment by comparing *trbd^C518A^* female (right) and *w^1118^* female (left) (p<0.05 and log2<-1 or >1). **(B)** Significant protein hits from ubiquitin GlyGly enrichment by comparing *trbd^C518A^* male (right) and *w^1118^* male (left) (p<0.05 and log2<-1 or >1). **(C)** Significant protein hits from ubiquitin GlyGly enrichment by comparing *trbd^C518A^* male (right) and *trbd^C518A^* female (left) (p<0.05 and log2<-1 or >1). **(D)** Significant protein hits from ubiquitin GlyGly enrichment by comparing *w^1118^* male (right) and *w^1118^* female (left) (p<0.05 and log2<-1 or >1). **(E)** Significant protein hits from total proteome by comparing *trbd^C518A^* female (right) and *w^1118^* female (left) (p<0.05 and log2<-1 or >1). **(F)** Significant protein hits from total proteome by comparing *trbd^C518A^* male (right) and *w^1118^* male (left) (p<0.05 and log2<-1 or >1). **(G)** Significant protein hits from total proteome by comparing *trbd^C518A^* male (right) and *trbd^C518A^* female (left) (p<0.05 and log2<-1 or >1). **(H)** Significant protein from total proteome by comparing *w^1118^* male (right) and *w^1118^* female (left) (p<0.05 and log2<-1 or >1).

**Figure S3. Ubiquitin linkage readings and quality control for proteomic experiments (A)** PCA analysis of ubiquitin GlyGly protein readings. **(B)** PCA analysis of total proteome readings. **(C)** Distribution of ubiquitin GlyGly protein readings. **(D)** Distribution analysis of total proteome readings. **(E)** No significant difference in Lys6, 11, 48 and 63 ubiquitin chain accumulation between *trbd*^C518A^ and *w*^1118^.

**Figure S4. Ubiquitylation event comparisons between genotypes. (A)** Ubiquitylation events when comparing *trbd^C518A^* female to *w^1118^* female. **(B)** Ubiquitylation events when comparing *trbd^C518A^* male to *w^1118^* male. **(C)** Ubiquitylation events when comparing *trbd^C518A^* male to *trbd^C518A^* female. **(D)** Ubiquitylation events when comparing *w^1118^* male to *w^1118^*female. Regulatory-related events refer to proteins only presented in the ubiquitome; expression-related events refer to proteins found in both ubiquitome and proteome; degradation/stabilisation-related events refer to proteins only found in the proteome.

**Table S1.** A list and GO data analysis for proteins with significant representation in the proteome and/or ubiquitinome (Table S1 contains the data for **Fig 5** and Fig S2).

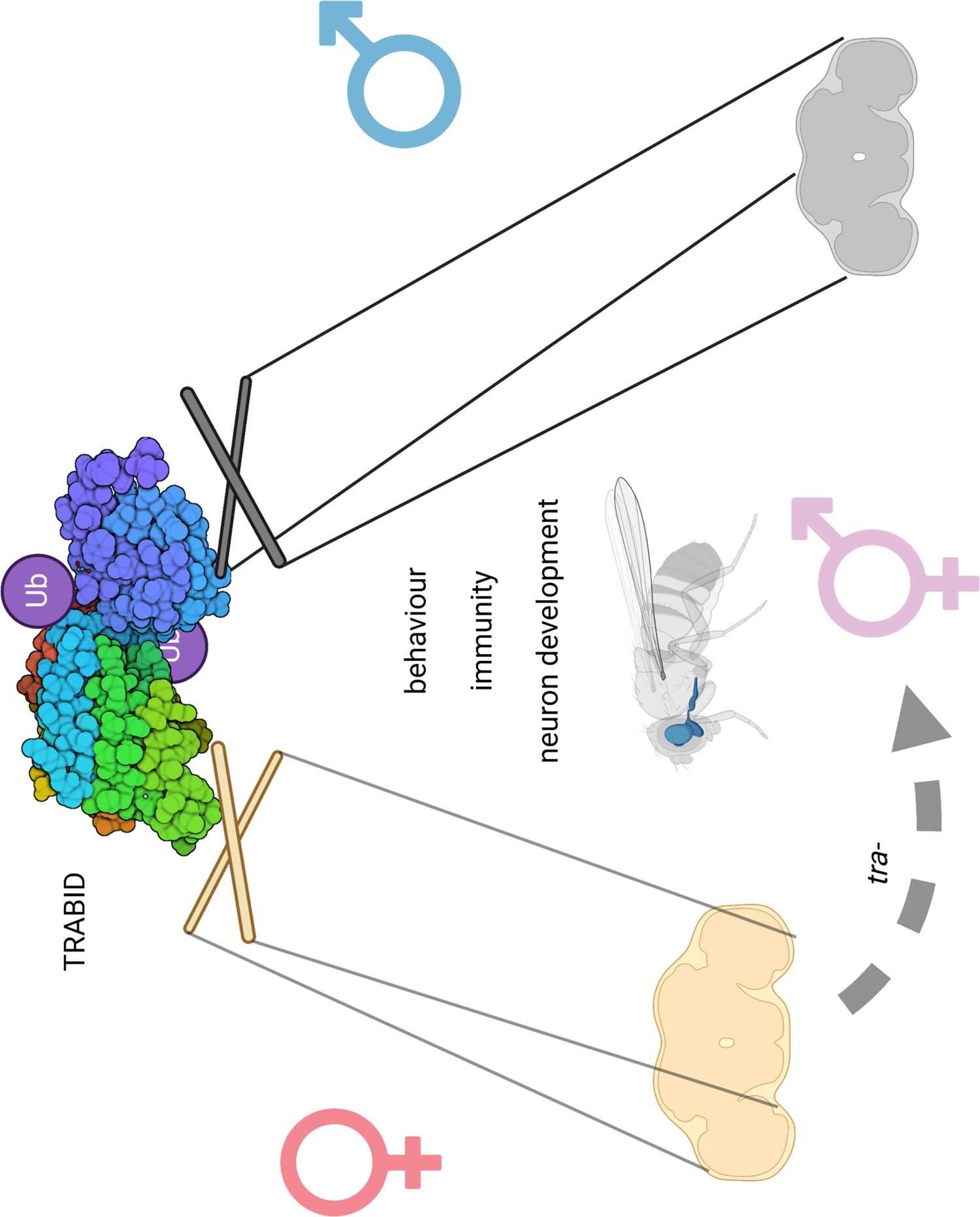

