## Supplementary figures and images for "Sex-specific deubiquitylation drives immune-related neurodegeneration in *Drosophila*"

### Fig S1

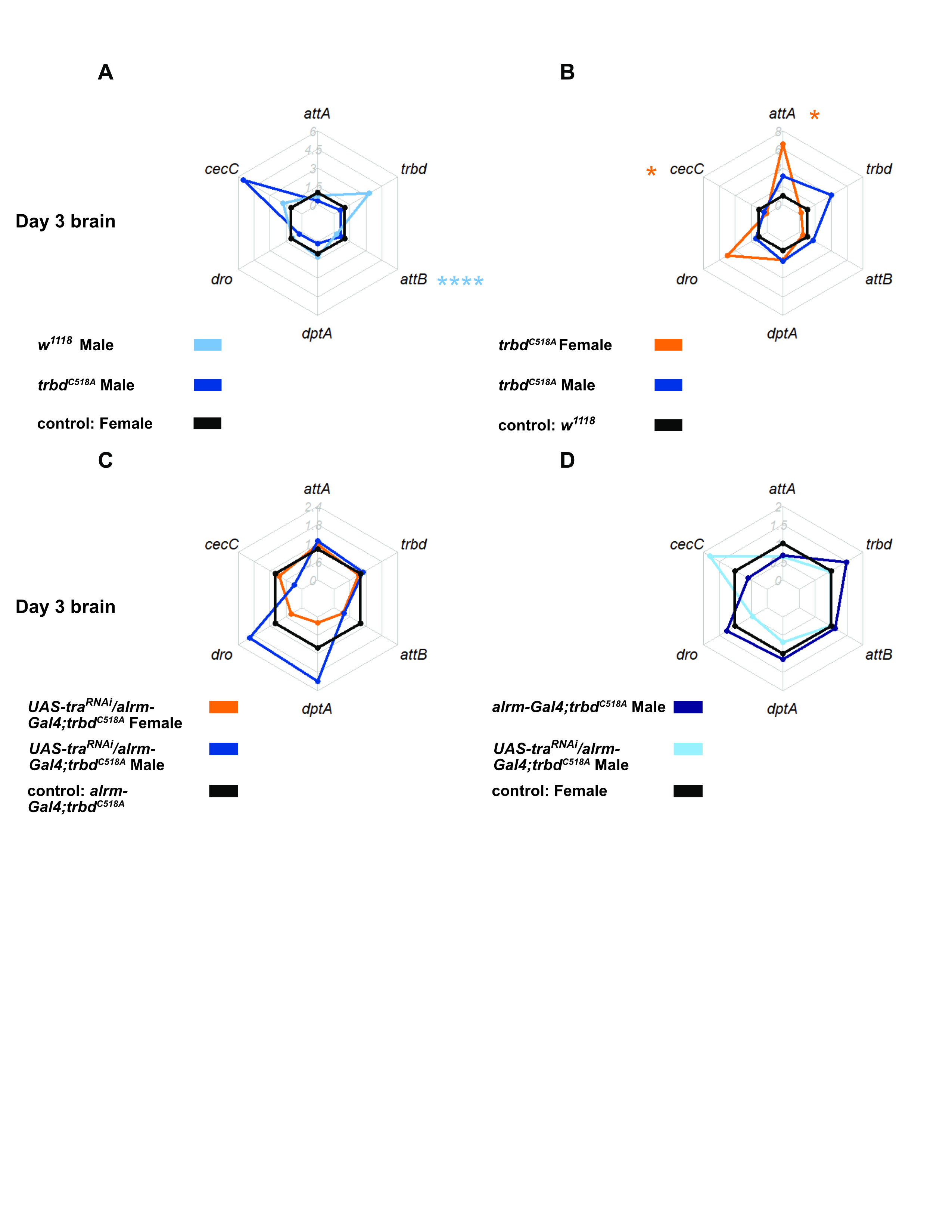

### Fig S2

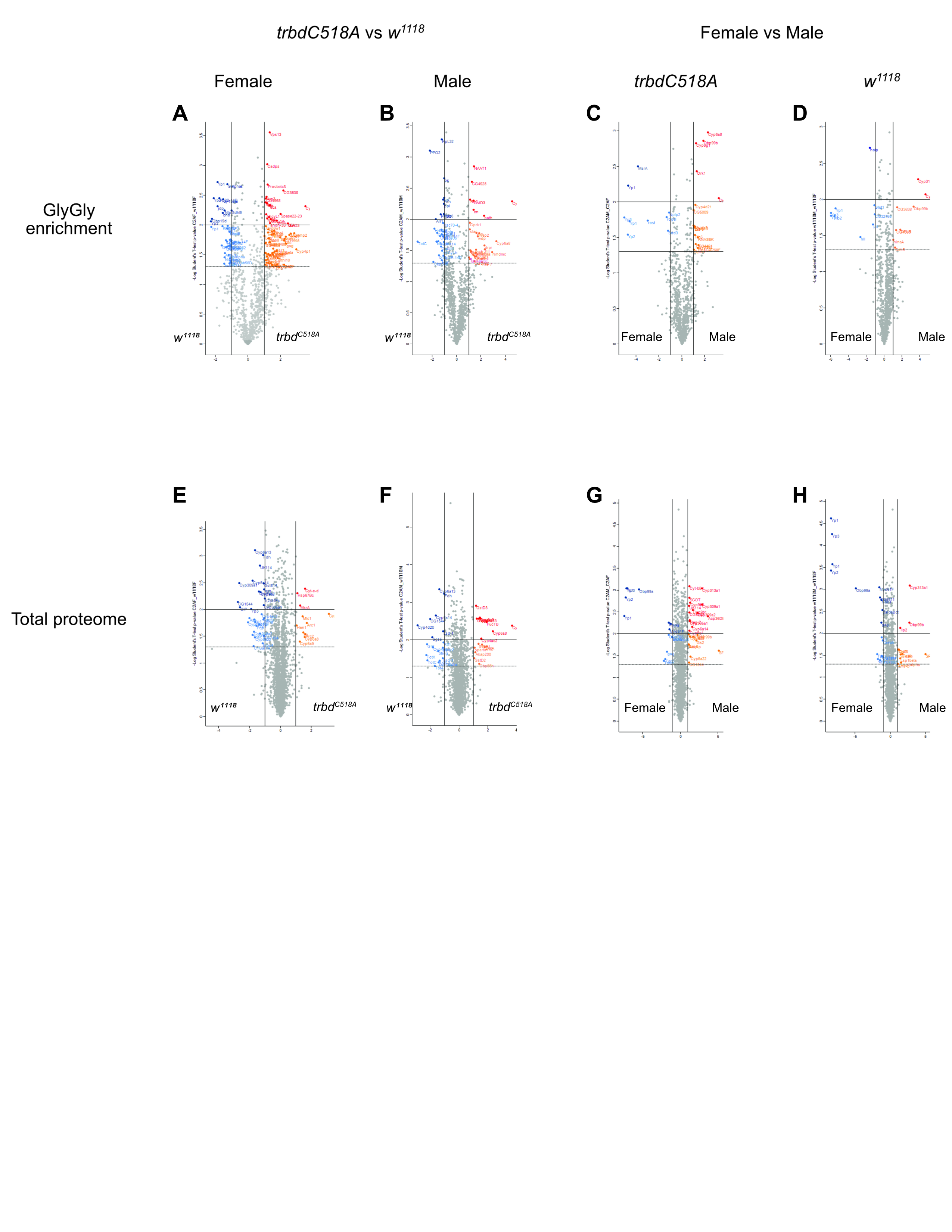

### Fig S3

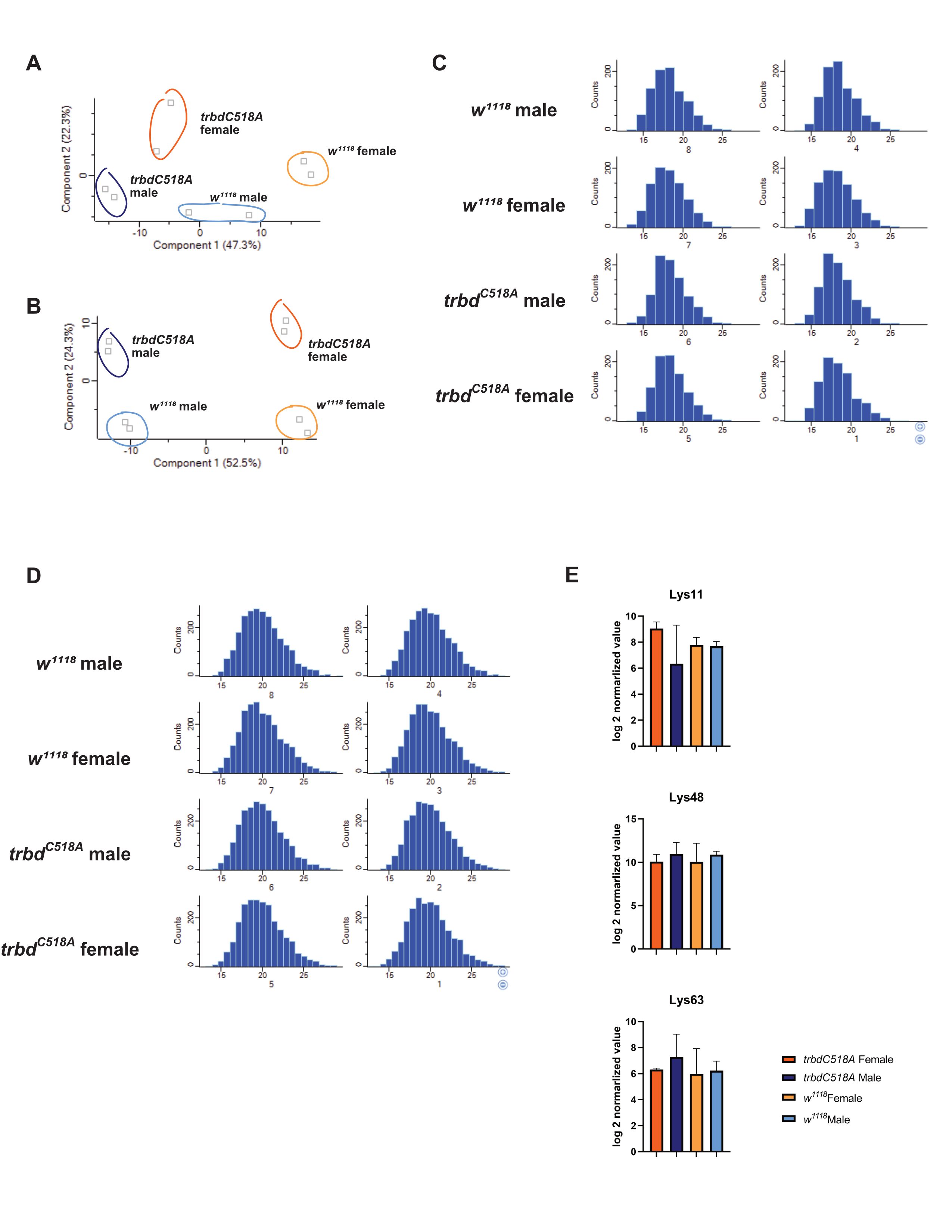

### Fig S4

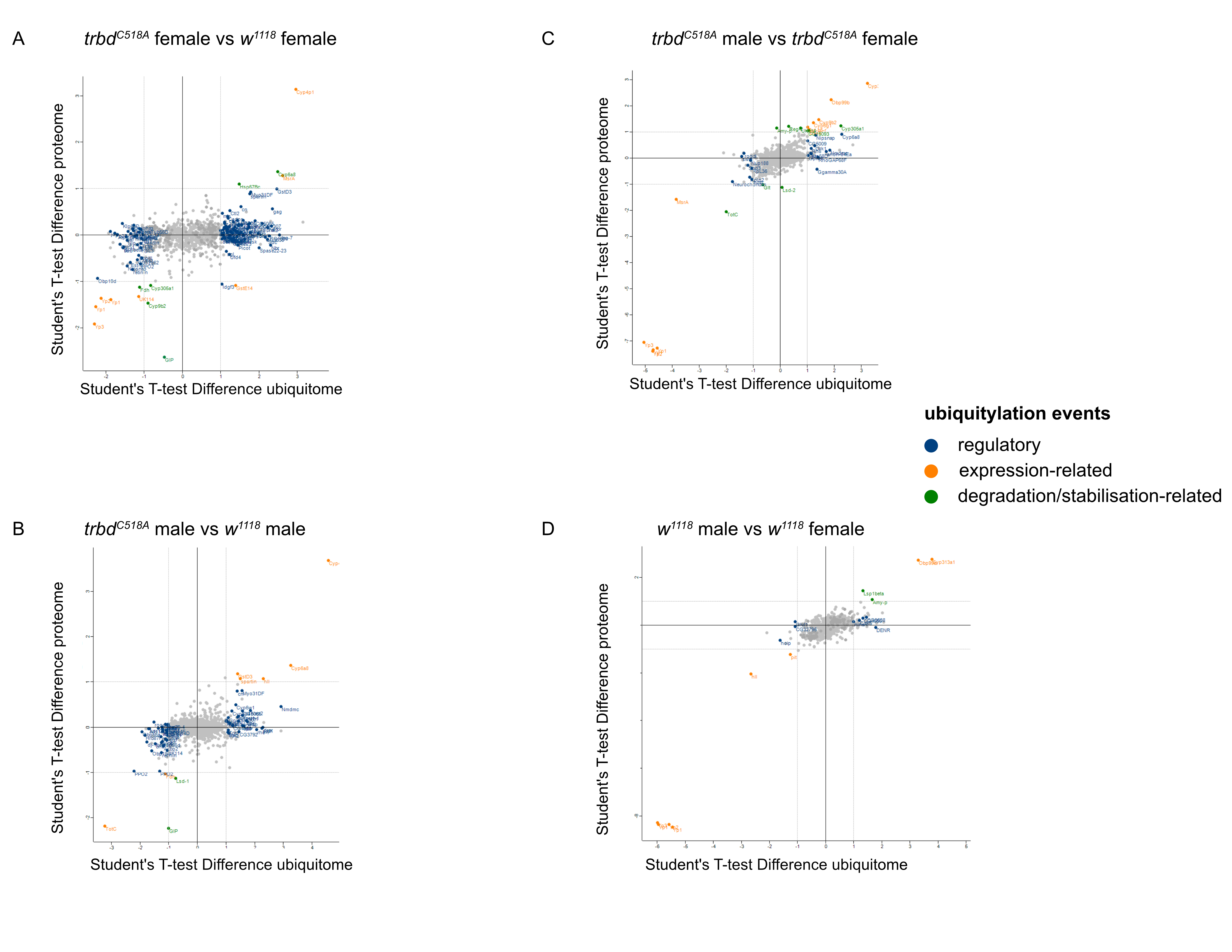
